# Notch-dependent DNA *cis*-regulatory elements and their dose-dependent control of *C. elegans* stem cell self-renewal

**DOI:** 10.1101/2021.11.09.467950

**Authors:** Tina R Lynch, Mingyu Xue, Cazza W. Czerniak, ChangHwan Lee, Judith Kimble

## Abstract

A long-standing biological question is how DNA *cis*-regulatory elements shape transcriptional patterns during metazoan development. The use of reporter constructs, cell culture and computational modeling has made enormous contributions to understanding this fundamental question, but analysis of regulatory elements in their natural developmental context is an essential but rarely used complement. Here, we edited Notch-dependent *cis*-regulatory elements in the endogenous *C. elegans sygl-1* gene, which encodes a key stem cell regulator. We then analyzed the *in vivo* consequences of those mutations – on both gene expression (nascent transcripts, mRNA, protein) and stem cell maintenance. Mutation of a single element in a three-element homotypic cluster reduced expression as well as stem cell pool size by about half, while mutation of two elements essentially abolished them. We find that LBS number and LBS neighborhood are both important to activity: elements on separate chromosomes function additively, while elements in the same cluster act synergistically. Our approach of precise CRISPR/Cas9 gene editing coupled with quantitation of both molecular and biological readouts establishes a powerful model for *in vivo* functional analyses of DNA *cis*-regulatory elements.

**Summary statement:** Notch-dependent DNA *cis*-regulatory elements work together in their developmental context to shape a transcriptional gradient, control stem cell pool size, and govern differentiation onset.

## Introduction

Cell signaling and transcriptional regulation are central to metazoan development. Though signaling pathways vary tremendously, a common theme is their patterned control of gene expression via DNA *cis*-regulatory elements (CREs). Traditional analyses of signaling rely on manipulations of the pathway’s core components (e.g. Austin and Kimble, 1987; Beumer and Clevers, 2021; Lee et al., 2016). Such pathway-level intervention is powerful, but likely impacts multiple target genes and circuits. An alternative approach manipulates signal-dependent CREs in the signaling target gene. Metazoan CREs have been subjected to many elegant analyses over decades, with integration of empirical data and computational modeling being particularly useful (e.g. Andersson and Sandelin, 2020; Bentovim et al., 2017; Ezer et al., 2014; Giorgetti et al., 2010; Lammers et al., 2020; Wong and Gunawardena, 2020). The advent of CRISPR/Cas9 gene editing has now made metazoan CREs accessible in their natural developmental context. The analysis of CREs in endogenous genes has the potential not only to solidify principles gleaned from the classic more artificial experiments but also to advance our understanding of how *cis*-regulatory elements function during development.

Here, we take advantage of a well-established model system to investigate how a cluster of CREs controls development *in vivo.* We focus on Notch-dependent CREs in the *C. elegans sygl-1* gene, which encodes a key regulator of self-renewal in germline stem cells (GSCs) (reviewed in Hubbard and Schedl, 2019). In this small nematode, GLP-1/Notch signaling from the niche activates transcription of two target genes, *sygl-1* and *lst-1* (Kershner et al., 2014; Lee et al., 2016; Shin et al., 2017) (Fig. 1A). These genes are functionally redundant: either *sygl-1* or *lst-1* can maintain GSCs on its own, but removal of both genes triggers premature differentiation and loss of GSCs, the Glp phenotype (Fig. 1B) (Kershner et al., 2014). Indeed, *sygl-1* and *lst-1* are likely the only Notch targets responsible for GSC maintenance (Chen et al., 2020). We previously reported that *sygl-1* and *lst-1* transcription is graded in germ cells within the niche (Fig. 1C) (Kershner et al., 2014; Lee et al., 2016), and that SYGL-1 and LST-1 proteins are restricted to a distal region within the progenitor zone (PZ), where GSCs reside (Fig. 1C, yellow) (Haupt et al., 2019; Shin et al., 2017). When SYGL-1 expression is expanded, the GSC pool also expands, suggesting that SYGL-1 spatial extent determines where GSC daughters transition from a stem cell state to one primed for differentiation (Shin et al., 2017). However, only *sygl-1* null mutants were available before this work, so the impact of reduced *sygl-1* was unknown.

**Figure 1.**
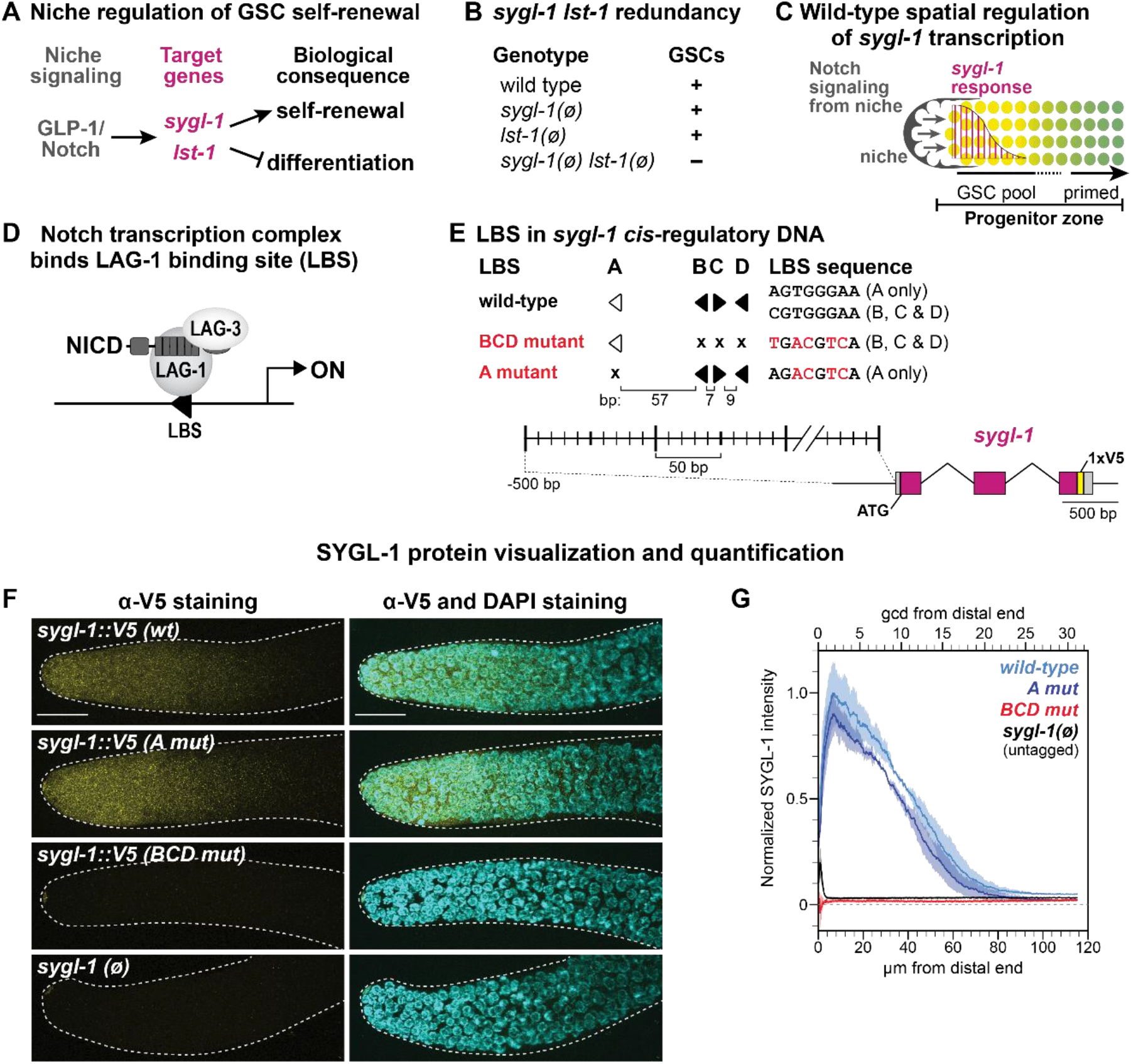
Identification of functional LBSs in *sygl-1 cis*-regulatory DNA. **A.** *C. elegans* germline stem cell (GSC) molecular regulators. **B.** Loss of either *sygl-1* or *lst-1* permits adult GSC self-renewal, but loss of both produces a Glp phenotype (small sterile germline, loss of GSCs). **C.** *C. elegans* distal gonad. The niche (gray) is a somatic cell at the distal end. GLP-1/Notch signaling (arrows) maintains a pool of GSCs in the distal progenitor zone (yellow) and activates graded *sygl-1* transcription (magenta). Germ cells in the proximal progenitor zone become primed for differentiation (green). **D.** Trimeric Notch transcriptional activation complex: Notch intracellular domain (NICD), Mastermind-like coactivator (LAG-3 in *C. elegans*), CSL DNA-binding member (LAG-1 in *C. elegans*). **E.** Filled arrowheads: canonical 5’ YGTGRGAA 3’ LBS; open arrowheads: noncanonical. Arrowheads point right for 5’ CGTGGGAA 3’ and left for its complement 5’ TTCCCACG 3’. Mutant LBS (x): 5’ TGACGTCA 3’ for LBS B, C, and D ; 5’ AGACGTCA 3’ for LBS A. Spacing in bp between each LBS is shown to scale. A 1xV5 epitope tag (yellow) was inserted just before the stop codon to visualize SYGL-1 protein. Exons, magenta; UTRs, gray. **F.** Representative maximum-intensity z-projections of V5-stained dissected distal gonads. Scale bar: 20 μm. Strain genotypes listed in Table S1. **G.** FIJI quantification of α-V5 signal normalized to *sygl-1::V5(wt)* (see Methods). Position measures are germ cell diameters (gcd) above and microns (μm) below. Total gonads scored from two independent experiments: *wt*, 29; *A mut*, 31; *BCD mut*, 25; *sygl-1(ø),* 24.

The *sygl-1* gene is well poised for CRE mutational analyses. Its transcription in GSCs relies on signaling by a single pathway, the Notch pathway (Kershner et al., 2014), and its wild-type transcriptional response to Notch signaling has been described at high resolution (Lee et al., 2016). Notch activates transcription via a highly conserved tripartite complex that includes LAG-1, a member of the CSL (CBF1-RBPJκ/Su(H)/LAG-1) family of DNA-binding proteins; the intracellular domain of the Notch receptor (NICD); and the LAG-3/SEL-8 Mastermind-like co-activator (Fig. 1D) (reviewed in Greenwald and Kovall, 2013). LAG-1 anchors the complex to DNA at LAG-1 binding sites (LBSs), motifs that are highly conserved for CSL proteins across phylogeny (Christensen et al., 1996; Nellesen et al., 1999; Tun et al., 1994). The *sygl-1* 5’ flanking region possesses four computationally predicted LBSs (Fig. 1E) (Yoo et al., 2004). Moreover, a 1 kb *sygl-1* DNA fragment harboring those LBSs drives GFP reporter expression in GSCs (Kershner et al., 2014). Wild-type *sygl-1* transcripts are graded across the GSC pool and become low or undetectable as GSCs are triggered to begin differentiation (Lee et al., 2016) (Fig. 1C). SYGL-1 protein, visualized with a V5 epitope tag, is patterned similarly to *sygl-1* RNA (Shin et al., 2017). Far more proximally in the germline, *sygl-1* transcription becomes independent of Notch signaling (Kershner et al., 2014; Lee et al., 2016). Previous quantitative analyses of Notch-dependent *sygl-1* transcription laid a critical foundation for our investigation of *sygl-1* CRE mutations. Single molecule fluorescence *in situ* hybridization (smFISH) in fixed gonads visualized nascent transcripts at nuclear active transcription sites (ATS) plus cytoplasmic mRNAs, and also showed that a weakened receptor forms a shallower gradient (Lee et al., 2016). Live imaging extended the smFISH results to reveal that the strength of Notch signaling corresponds to the duration of a *sygl-1* transcriptional burst (Lee et al., 2019), a result confirmed by a parallel study in Drosophila embryos (Falo-Sanjuan et al., 2019).

Our approach to *in vivo* investigations of CRE function takes advantage of a metazoan exemplary for its relative simplicity and tractability. Multiple and diverse CREs have been implicated in regulation of transcriptional patterns during development, which makes CRE mutations difficult to interpret (Kuang et al., 2021; Li et al., 2021; Park et al., 2019). However, tackling CRE function in a well-defined model like *C. elegans*, and the *sygl-1* gene in particular, strips layers of complexity. For example, in flies and mammals, a cooperative physical interaction between neighboring NICD proteins promotes transcriptional synergy when CSL binding sites are spaced ~15-17 bp apart and arranged with head-to-head polarity (Arnett et al., 2010; Bailey and Posakony, 1995; Cave et al., 2005; Kobia et al., 2020; Kuang et al., 2021; Nam et al., 2007; Nellesen et al., 1999; Severson et al., 2017). By contrast, the molecular interface responsible for that NICD interaction is not conserved in nematodes, and LBS polarity is apparently not critical for Notch-dependent expression in *C. elegans* (Nam et al., 2007; Neves et al., 2007). The lack of this cooperative NICD interaction in nematodes removes one layer of complexity and so has potential to reveal principles relevant to homotypic clusters more broadly.

Here, we couple Cas9 gene editing with quantitative *in vivo* analyses to investigate the functions of individual LBSs within a homotypic cluster. We find that LBS number emerges as a major force in shaping the developmental gradient. The LBS dose modulates both probability and intensity of *sygl-1* transcription, abundance of *sygl-1* mRNA and protein, and size of the GSC pool. Furthermore, we discover that LBSs on separate chromosomes act additively, while LBSs on the same chromosome act synergistically. Finally, we identify rough boundaries for the functional threshold of SYGL-1 abundance that is required for self-renewal. This *in vivo* investigation provides a model approach for learning how DNA *cis*-regulatory elements transform signaling inputs into reproducible patterns of gene expression during development.

## Results

### A cluster of three *sygl-1* LBSs activate Notch-dependent expression

Four computationally predicted Notch-dependent *cis*-regulatory elements are named LBS A through D (Fig. 1E). LBS B, C and D exist in a cluster while LBS A resides upstream of the cluster. In other *Caenorhabditids, sygl-1* 5’ flanking regions contain LBS clusters in species-specific patterns containing at least two LBSs (Fig. S1A). LBS B, C, and D sequences adhere to the canonical CSL binding motif, while LBS A lacks the initial pyrimidine and is thus noncanonical. Such noncanonical LBSs compete poorly for CSL binding in gel shift assays (Christensen et al., 1996; Nellesen et al., 1999; Torella et al., 2014; Tun et al., 1994), and are expected to be weaker sites *in vivo*.

We hypothesized that the BCD cluster is largely responsible for Notch-dependent regulation of *sygl-1* expression in GSCs. To test this idea, we used Cas9 gene editing to generate two mutants in endogenous *sygl-1* DNA: *A mut* changes 4 basepairs in LBS A, and *BCD mut* changes 5 bp in each of LBS B, C, and D (Fig 1E, see Methods mutant design). Both *A mut* and *BCD mut* animals were fertile and their germlines were of normal size and organization in the presence of wild-type *lst-1*, the functionally redundant counterpart of *sygl-1.* However, when *lst-1* was removed, *BCD mut* adults were sterile with a Glp phenotype (loss of GSCs), while *A mut* adults remained fertile with a normal germline. Therefore, *BCD mut* had a dramatic effect on germline development, but *A mut* had no apparent effect. To visualize SYGL-1 protein, we inserted a V5 epitope tag in each mutant (Fig 1E). Initial analyses focused on the distal gonad, where *sygl-1* expression is Notch-dependent, and were performed in animals with wild-type *lst-1* to ensure a healthy germline. *A mut* had a wild-type pattern of SYGL-1 protein, both in abundance and distribution, but *BCD mut* had no SYGL-1 (Fig. 1F-G). Consistent with the idea that LBSs drive Notch-specific *sygl-1* transcription, the Notch-independent production of SYGL-1 appeared comparable in the proximal germlines of wild-type, *A mut*, and *BCD mut* (Fig. S1B). In an attempt to increase expression, we transformed LBS A to a canonical motif, but found no effect (Fig. S1C-D). We conclude that the BCD cluster is responsible for Notch regulation of SYGL-1 expression in GSCs.

To assay functions of individual elements in the BCD cluster, we mutated the sequence of each LBS from the canonical 5’ CGTGGGAA 3’ motif to 5’ TGACGTCA 3’ (differences from canonical motif underlined; see Methods). The three LBSs were mutated singly and in all possible pairs (Fig 2A). We then subjected all mutants to a series of molecular and biological assays (Fig 2-5). As a control for the sequence change, we created a distinct LBS D mutant, *alt D mut*, which had effects comparable to *D mut* (Fig. S5). The *sygl-1* LBS single and double mutants were all homozygous fertile in the presence of wild-type *lst-1*, the functionally redundant counterpart of *sygl-1* (Fig. 1A-B). Molecular assays were therefore done with *lst-1(+)* to ensure a healthy germline, but stem cell assays were done in an *lst-1(ø)* background. The following sections describe molecular assays first and then biological assays.

**Figure 2:**
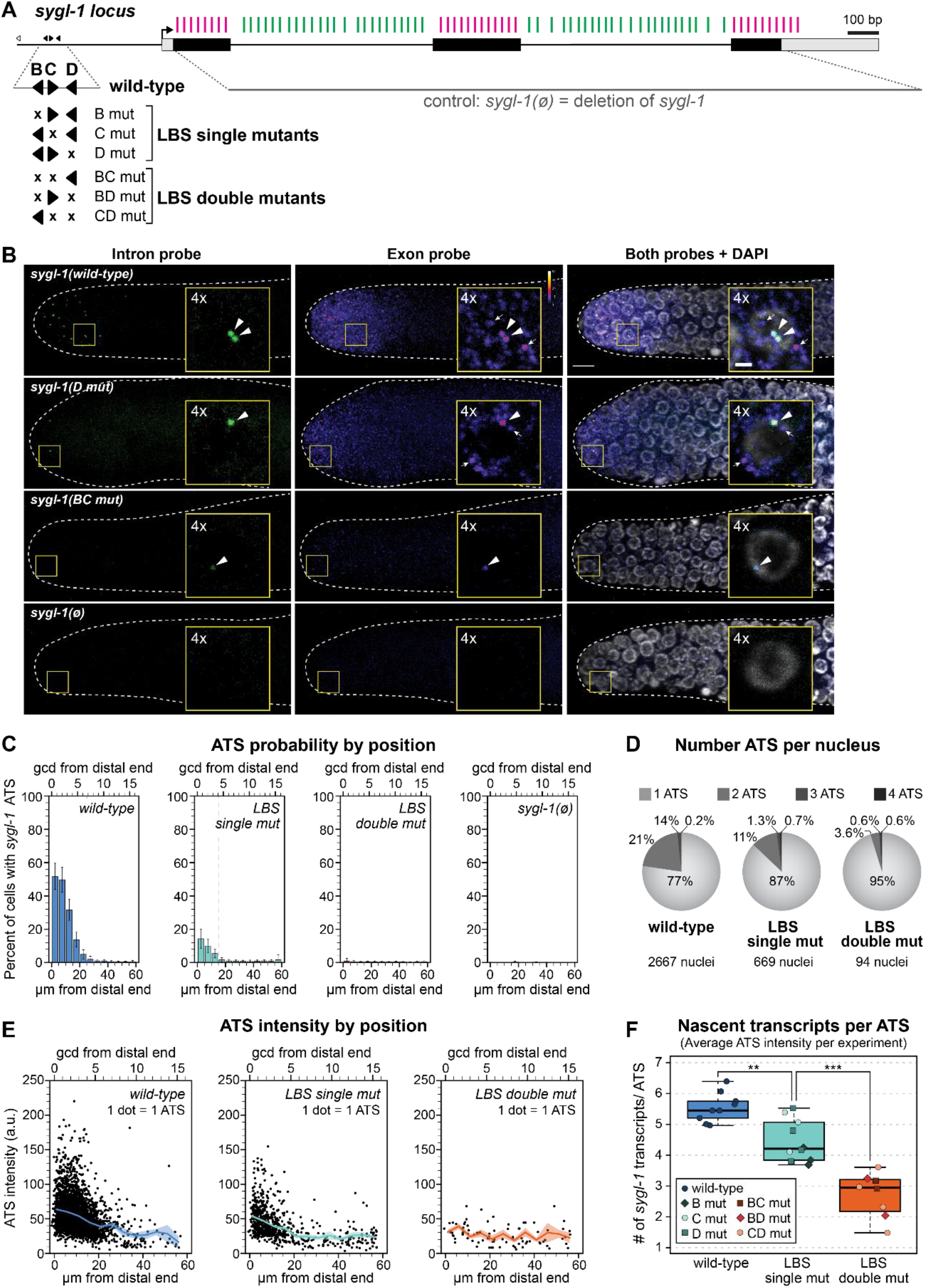
LBS mutations weaken niche-dependent transcriptional response of *sygl-1*. **A.** Black boxes: exons; gray boxes: untranslated regions; black arrow: predicted transcription start site; smFISH probes to exons (magenta) and introns (green) (see Methods). The *sygl-1(ø)* control removes *sygl-1* sequence that probes detect. Expansion shows individual LBSs and their mutations; conventions as in Fig1E. LBS mutants in this figure are not V5-tagged. **B.** Representative images of *sygl-1* smFISH in distal end of dissected gonads. Exon channel colors reflect intensity to show mRNA without saturating ATS (see Methods; color key in upper right; pixel intensities 3-50). DAPI shown in grayscale. Main images: maximum-intensity z-projections; insets: 4x-magnified maximum-intensity projections (four 0.3 μm slices). Arrowheads: ATS (overlapping intron/exon/DAPI). Arrows: cytoplasmic mRNA (exon only). Strain genotypes listed in Table S1. **C-F.** See Fig S2E for details (e.g. total # gonads scored. **C.** Percentage of nuclei with ≥1 ATS as a function of distance (units as in Fig 1G). Extent of transcription indicated by gray dashed line (where <5% cells contain ≥1 ATS). Error bars: SEM. **D.** Numbers in pie charts are averages between experiments (see Methods). Nuclei with zero *sygl-1* ATS excluded. **E.** Each dot represents one ATS (see Methods); see Fig S2D for details (e.g. total ATS scored). Solid line: mean (5 μm intervals); shaded: SEM. **F.** ATS intensities ÷ 10 roughly estimate # primary transcripts/ATS (see Methods). Boxplot center lines: median (wt: 5.1; singles; 3.7; doubles: 2.6); see Methods for BoxPlotR conventions. Each dot represents mean of all ATS/experiment (irrespective of position). Total # experiments: wt: 9; singles: 10; doubles: 8. Experiments with zero ATS detected (*BC mut*, *BD mut*) are not represented.

### *sygl-1* LBS mutants fire fewer and weaker active transcription sites

We visualized *sygl-1* RNAs *in situ* with high resolution and spatiotemporal precision using single molecule FISH (smFISH). Fixed gonads were treated with two probe sets that were distinctly labeled to *sygl-1* exons or introns (Fig. 2A). Figure 2B shows representative smFISH images for LBS single and double mutants plus wild-type and *sygl-1(ø)* controls. The *sygl-1* intron probe detected nascent transcripts in the nucleus as bright spots (Fig. 2B, green, left column), while the *sygl-1* exon probe detected both nuclear bright spots and a multitude of dim spots in the cytoplasm (Fig. 2B, middle column, color scaled by intensity). Overlapping exon and intron probe signals in the nucleus identified *sygl-1* active transcription sites (ATS) (Fig. 2B arrowheads), while the dim cytoplasmic spots identified *sygl-1* mRNAs (Fig. 2B, arrows) (see Lee et al. 2016 for additional *sygl-1* ATS and mRNA validation).

We quantified effects of LBS mutations on transcription in 3D with a MATLAB code used previously to score wild-type *sygl-1* transcription (Crittenden et al., 2019; Lee et al., 2016). As established in those earlier works, the percentage of cells with any *sygl-1* ATS provides a measure of *sygl-1* transcriptional probability, and the intensity of each ATS signal provides a measure of firing strength. When LBS mutants were scored individually, each single mutant lowered transcription substantially, and each double mutant nearly abolished it (Fig. S2). Because the three LBS single mutants all had similar effects (Fig. S2B,D-E), as did the three double mutants (Fig. S2C,D-E), and because all made far fewer ATS than wild-type (Fig. S2B-E), we pooled ATS data into collective “LBS single mut” and “LBS double mut” datasets (Fig. 2C-F) for further analyses.

Transcriptional probability was scored in cells as a function of distance from the distal end of the gonad. For wild-type, probability was highest at the distal end adjacent to the niche and lowered progressively with distance from the end (Fig. 2C), as previously reported (Crittenden et al., 2019; Lee et al., 2016). In LBS single mutants, the probability was lower than wild-type, but similarly graded; in LBS double mutants, the probability was near zero (Fig. 2C, Fig. S2A-C). Overall, transcriptional probabilities were dramatically attenuated in both height and spatial extent along the gonadal axis with LBS double mutants having the most severe effect. As a complementary measure of transcriptional probability, we scored the number of *sygl-1* ATS in each nucleus. Because *sygl-1* transcriptional probability is stochastic and unrelated to cell cycle stage (Lee et al., 2016), any nucleus in the dividing pool of germ cells might possess zero to four *sygl-1* ATS, depending on chromosome replication and probability of *sygl-1* transcription. Consistent with our first measure, the percentage of nuclei with more than one ATS was lower in LBS single mutants than wild-type and even lower in LBS double mutants (Fig. 2D, S2E). Thus, LBS dose regulates the probability of Notch-dependent *sygl-1* transcription.

We next scored ATS signal intensity as a metric of transcriptional firing strength. Intensity values from individual ATS were captured in the exon channel and normalized using the average raw intensity of cytoplasmic mRNA (see Methods). When plotted as a function of position along the gonadal axis, mean ATS intensities appeared to decrease further from the distal end (Fig. 2E). To estimate the average number of nascent transcripts at each ATS, we divided the average normalized ATS intensities by the average normalized mRNA intensity (see Methods). By this measure, an ATS in LBS single mutants generated ~20% fewer nascent transcripts than wild-type, on average, and an ATS in LBS double mutants made ~50% fewer transcripts, on average (Fig. 2F). We conclude that *sygl-1* LBS dose regulates both the probability and intensity of *sygl-1* transcription.

### *sygl-1* LBS single mutants reduce *sygl-1* mRNA and protein subtantially

We analyzed *sygl-1* mRNAs and SYGL-1 protein in all three LBS single mutants to learn how their reduced transcription affects molecular output in the distal gonad. The *sygl-1* mRNAs were quantified from the same smFISH images used for ATS analyses, where LBS mutants were not epitope-tagged. SYGL-1 protein was quantified with immunostaining of a V5 epitope tag inserted at the SYGL-1 C-terminus of each LBS mutant. This V5 tag has no detectable effect on SYGL-1 protein function (Shin et al., 2017).

To quantitate *sygl-1* mRNAs, we used MATLAB to detect and count cytoplasmic spots in the exon channel of smFISH images, as described previously (Crittenden et al., 2019; Lee et al., 2016) (see Methods, Fig. S3A). In LBS single mutants, mRNA abundance was reduced to slightly less than half of wild-type (Fig. 3A,C), and spatial extents along the gonadal axis were shorter than wild-type by about 5 μm or 1-2 cell rows (Fig. 3A, vertical lines).

**Figure 3:**
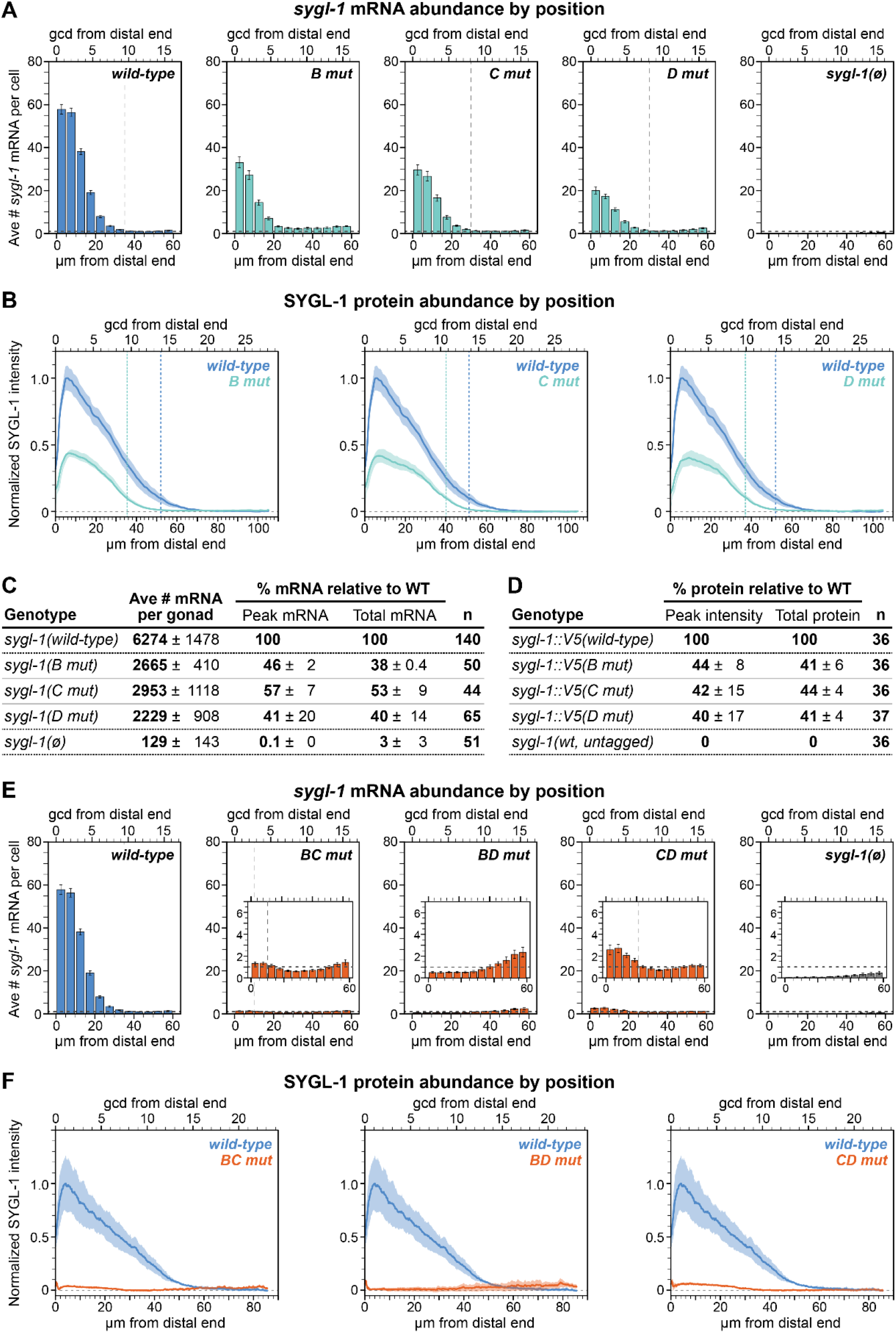
Molecular *sygl-1* output is reduced. RNA data from smFISH experiments (Fig. A, C, E) and protein data from α-V5 staining (Fig. B, D, F) were both measured in an *lst-1(+)* background. Strain genotypes listed in Table S1; all data is from adult animals. **A-D.** LBS single mut *sygl-1* mRNA and protein are reduced to approximately half of wild-type. **E-F.** LBS double mut *sygl-1* mRNA and protein are reduced to near zero. In Fig. 3E, insets are used to better visualize where mRNA is above or below background. **A,E.** Average # *sygl-1* mRNA/cell (see Methods). Position measures as in Fig 1G. Error bars: SEM. Vertical dashed lines: spatial extent, where values fall below background and/or reach minimum. Horizontal dashed lines: background (from *sygl-1(ø),* see Methods). Total # gonads scored: see Fig S2E. **B,F.** Quantification of α-V5 immunofluorescence (see Methods). Solid lines: mean; shading: SEM. y axis “0” (dashed line) represents values from untagged control. Small peaks 0-3 μm from distal end are nonspecific V5 signal (see Methods). Dashed lines: spatial extent, where values fall below 10% WT maximum. Total # gonads scored in Fig. 3B listed in Fig3D; total gonads scored in Fig 3F from 3-4 experiments: *wt*: 61; *BC mut:* 49; *BD mut:* 59 *CD mut:* 39. **C.** Summary of mRNA data in LBS single mutants. Numbers are mean per experiment ± standard deviation between experiments (see Methods). n: # gonads scored in ≥3 experiments. Peak: mRNA/cell values from the 0-5 μm region as a percent of wild-type; total: numbers of mRNA per gonad as a percent of wild-type. **D.** Summary of protein data in LBS single mutants. Numbers are mean percentage of wild-type for each replicate ± standard deviation between replicates (see Methods). n: total gonads scored in 3 replicates.

To quantitate SYGL-1 protein, we stained LBS single mutants with α-V5 antibodies (Fig. S3B) and scored fluorescence intensities along the gonadal axis using FIJI (see Methods). In LBS single mutants, the peaks of the SYGL-1 gradients were reduced to about 40% of wild-type (Fig. 3B,D), and spatial extents were shorter by about 25% of wild-type (~15 μm or 3-5 cell rows) (Fig 3B, vertical lines). We conclude that the gradients of both *sygl-1* RNA and SYGL-1 protein are reduced to a similar degree in all three LBS single mutants.

### *sygl-1* LBS double mutants nearly eliminate *sygl-1* mRNA and protein

We also quantified *sygl-1* mRNA and SYGL-1 protein in LBS double mutants, using the same strategy as for LBS single mutants. The Notch-dependent *sygl-1* mRNA abundance was below background in the distal-most germ cells in *BD mut* and just above background in *BC mut* and *CD mut* (Fig. 3E, insets) (see Methods). SYGL-1 protein was also very low, but again only *BD mut* lacked distal Notch-dependent signal above background (Fig. 3F, horizontal dashed lines; Fig. S3D). We conclude that protein levels parallel mRNA levels in each of the three *sygl-1* LBS double mutants, which are expressed at or near zero.

### LBS single but not double mutants make enough SYGL-1 to maintain adult GSCs

The molecular quantitation described above was done in an *lst-1(+)* background to ensure a healthy germline. However, LST-1 masks stem cell defects in *sygl-1* LBS mutants, because of redundancy (Fig. 1B). To score effects of the LBS mutants on stem cell maintenance, we removed LST-1 genetically by introducing *lst-1(ø)* into each mutant. All three LBS single mutants were fertile in an *lst-1(ø)* background and had a germ line of normal size and organization. By contrast, all three LBS double mutants were sterile with tiny sperm-filled germlines, the Glp phenotype (Fig. 4A). Therefore, LBS single mutants generate SYGL-1 at an abundance above the functional threshold required to maintain adult GSCs, but LBS double mutants do not.

**Figure 4:**
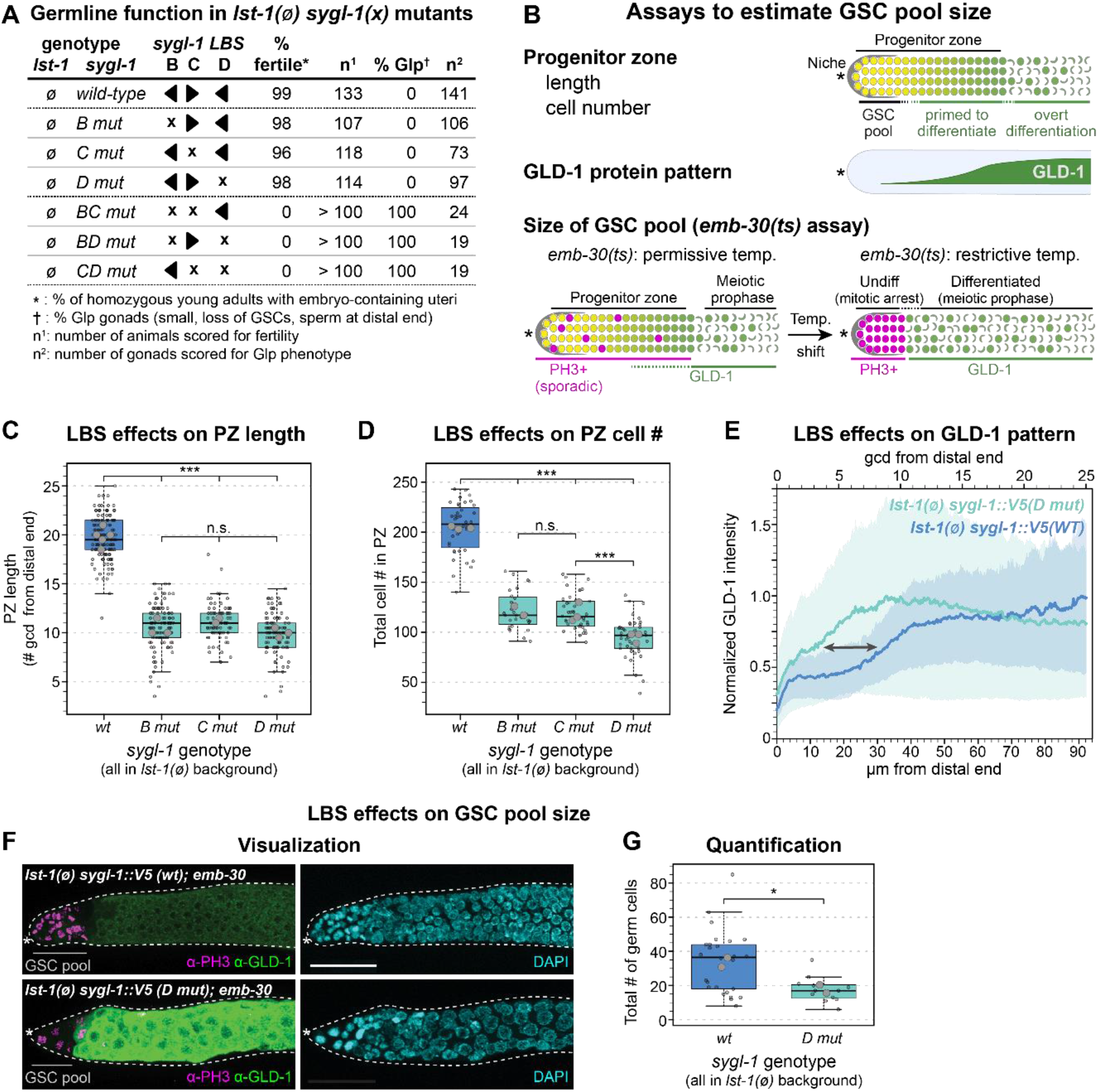
Effect of SYGL-1 abundance on GSC pool size. **A.** Percent of fertile animals was scored at low magnification and percent Glp animals scored at high magnification. The same gonads scored for %Glp (Fig4A) were used to score PZ length (Fig4C). **B.** Top, Progenitor Zone (PZ) includes distal GSC pool and GSC daughters primed to differentiate. Overt differentiation: visible meiotic prophase. Middle, increased GLD-1 protein reflects differentiation. Bottom, GSC pool size estimation; temperature-sensitive (ts). Left: at 15°C, *lst-1(ø) sygl-1(x); emb-30(ts)* PZ contains scattered M-phase (PH3+, magenta) cells and gradual increase in GLD-1 (green). Right: when shifted to 25°C, PZ germ cells arrest, defining two populations, mitotically-arrested distal PH3+ cells (inferred to be GSCs) and proximal GLD-1+, PH3-negative cells (inferred to be GSC daughters). *: distal end. **A, C-G.** Strain genotypes in Table S1. **C-D.** Boxplot conventions match Fig 2F. Small translucent data points: individual gonads; larger data points: replicate averages. Data collected from each gonad were fitted to a linear mixed effects model; Tukey’s post-hoc test was used to make pairwise comparisons between genotypes. ***: p < 0.0001; n.s., not significant (p ≥ 0.01). **C.** Average # germ cell diameters (gcd) in PZ were counted by eye. Total gonads scored in 2-5 experiments: *wt*: 141; *B mut:* 106; *C mut:* 73; *D mut:* 97. **D.** Total # cells in PZ were counted with Imaris software (see Methods). Total gonads scored in 3 independent experiments: *wt*: 40; *B mut:* 28; *C mut:* 42; *D mut:* 42. **E.** Fixed adult gonads were stained with polyclonal antibody to GLD-1 (gift from T. Schedl). Gray arrow: distal shift of GLD-1 accumulation. Position measures as in Fig. 1G. Signal intensities normalized against internal controls (see Methods). Solid lines: mean; shading: SEM. 24 gonads/genotype scored from 2 replicates. **F.** Representative maximum-intensity projections. Left, α-PH3 (magenta), α-GLD-1 (green). Gray line: relative lengths of GSC pools. Right, DAPI (cyan); scale bar: 20 μm. **G.** Estimated # GSCs in naïve pool; GLD-1-negative PH3+ cells were manually counted in FIJI. Boxplot conventions are as in Fig 2F, 5C-D. Total gonads scored in 2 experiments: *wt*: 26; *D mut:* 12. Meiotic gonads excluded from *D mut* (see Methods). *: p = 0.05. Student’s two-tailed T-test conducted on replicate averages; homoscedasticity assumed.

### GSC pool size is reduced in LBS single mutants

Although the LBS single mutants maintain adult GSCs, we suspected that their lower SYGL-1 abundance might maintain fewer GSCs and thus shrink GSC pool size. To test this idea, we used three complementary assays (Fig. 4B). All were performed in an *lst-1(ø)* background to eliminate LST-1 redundancy. First, we measured progenitor zone (PZ) size, a common proxy for GSC pool size. Both PZ length along the gonadal axis and total PZ cell number were reduced by about half in all three LBS single mutants compared to wild-type (Fig. 4C-D). Second, we measured distribution of GLD-1, a marker of germ cell differentiation. In *lst-1(ø)* control gonads, GLD-1 abundance increased as germ cells move proximally through the progenitor zone (Fig 4E), as previously reported (Brenner and Schedl, 2016; Hansen et al., 2004; Jones et al., 1996). We focused on *D mut* for this and next assay, because the LBS single mutants had behaved similarly overall. In *D mut* gonads the GLD-1 increase shifted distally compared to the control (Fig. 4E, gray arrow). The likely interpretation of the shorter PZs and distally-shifted GLD-1 increase is that germ cells are triggered to begin differentiation more distally in LBS single mutants than controls. Third, we estimated GSC pool size, using the *emb-30* assay. This assay provides a rough but more direct measure than other assays (Fig. 4B, see legend for explanation). When shifted to restrictive temperature, *emb-30* mutants reveal a distal GSC pool and a proximal pool of GSC daughters starting to differentiate (Cinquin et al., 2010). To estimate GSC pool size in an LBS single mutant, we shifted adults to restrictive temperature for 12.5 hours, stained gonads with α-GLD-1 plus a marker of mitosis (α-PH3), and counted undifferentiated GSCs in the distal gonad (see Methods). *D mut* gonads had a visibly smaller GSC pool than the control (Fig. 4F), with 18 cells on average compared to 34 (Fig. 4G). Together, these assays provide complementary lines of evidence that LBS single mutants maintain a smaller-than-normal GSC pool.

### LBSs have roughly equivalent but not identical activivities

Data in Figures 2-4 support the idea that the three LBS in the *sygl-1* cluster have essentially equivalent roles in regulating *sygl-1* expression and GSC maintenance. Two clues that they might not be identical were that *BC mut* and *CD mut* made marginally more *sygl-1* mRNA and protein than *BD mut* (Fig 3E,F) and that *D mut* had fewer cells in the PZ than *B mut* or *C mut* (Fig. 4D). To investigate these differences in more depth, we turned to LBS double mutants: each leaves one LBS intact and can be used to score activity of that “solo” LBS.

To ask if the minor molecular differences among LBS double mutants lead to biological differences, we first measured PZ lengths. LBS double mutants do not have a PZ in an *lst-1(ø)* background, so we used an *lst-1(+)* background for this experiment. LBS double mutants had a shorter-than-normal PZ, similar to *sygl-1(ø)* and consistent with their near-complete elimination of SYGL-1 (Fig. 5A). However, PZ length was marginally shorter in *BD mut* than the other double mutants (p = 0.04), which were not different from each other (p = 0.10). Because this difference was so slight, we tried another approach. We removed *lst-1* genetically and counted total number of germ cells in fourth larval stage (L4) larvae. In control *lst-1(ø) sygl-1(ø)* larvae, GSCs begin differentiating in L1 stage (Kershner et al., 2014). Therefore, if L4 larvae have more germ cells than the *lst-1(ø) sygl-1(ø)* control, the solo LBS must have made more SYGL-1 than the null. Germ cells were counted with DAPI and a sperm marker in whole-fixed L4 larvae (see Methods). *BD mut* made the same number of germ cells as the control, but the other two LBS double mutants had five-fold more germ cells than the control (Fig. 5B). Therefore, solo LBS B and solo LBS D must each have weak transcriptional activity, at least in larvae, but solo LBS C is equivalent to the null.

**Figure 5:**
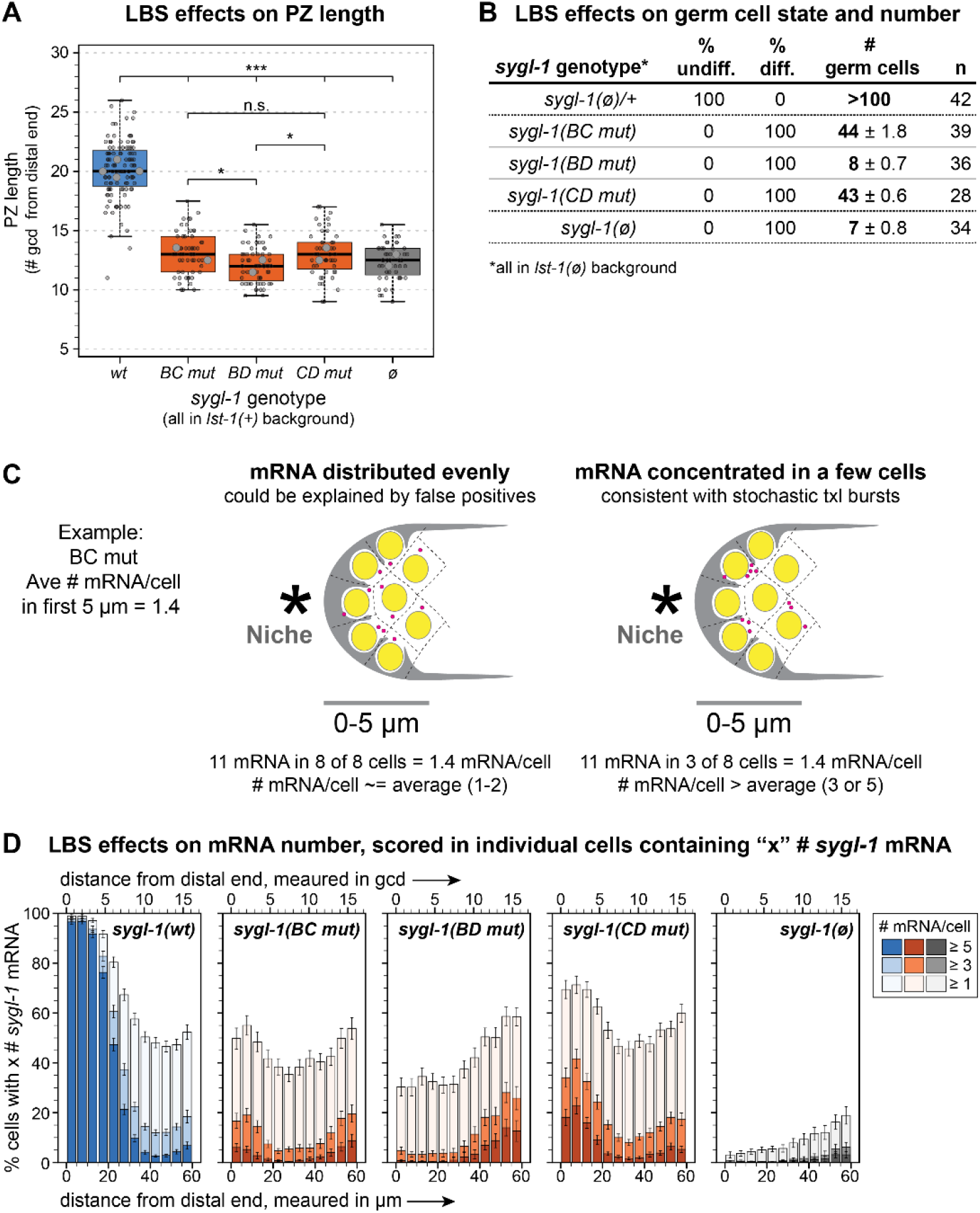
LBS double mutants probe individual LBS equivalence. **A-C.** Strain genotypes in Table S1. **A.** Boxplot conventions match Fig 4C-D. Total gonads scored in 2-4 experiments: *wt*: 111; *BC mut:* 57; *BD mut:* 59; *CD mut:* 48; *sygl-1(ø):* 48. Linear mixed effects model and Tukey’s post-hoc used as in Fig4C-D. *: p < 0.05 (p = 0.04 in both cases); *** p < 0.0001. All other pairwise comparisons are not significant (n.s.: p > 0.05). Not all n.s. comparisons shown. **B.** Total # germ cells (both gonad arms); average/replicate ± standard deviation between replicates. Percent of animals were scored as undifferentiated (undiff) or differentiated (diff) by their distalmost germ cells using DAPI morphology and/or SP56+ staining, and germ cell numbers were extrapolated (see Methods). n: total gonads scored in 2-4 experiments. Control was JK6401. **C.** Two explanations of an average of 1.4 mRNA/cell in a troop of ~8 cells (see Fig S4A). Left: 1-2 mRNAs in all cells; Right: no RNAs in some cells and 3-5 mRNAs in others. **D.** Percentage of total nuclei that contain ≥1, ≥3, or ≥5 *sygl-1* mRNA in each bin of distance from the distal end (Methods). Position measures as in Fig 1G. Error bars: SEM. Strains in *lst-1(+)* background.

The detectable biological activities of *BC mut* and *CD mut* made us wonder if the mRNA numbers in Figure 3E, which are averages, might be misleading. A previous work showed that adjacent germ cells can differ dramatically in mRNA number (Lee et al., 2016). Perhaps one or a few cells in LBS double mutants make considerably more than the average and others make none (Fig. 5C). To test this idea, we reassessed *sygl-1* RNA numbers on a cell-by-cell basis in the distal bin (0-5 μm) of wild-type, LBS single mutants, LBS double mutants and *sygl-1(ø).* Virtually all cells had ≥5 mRNAs in wild-type and most cells had ≥5 mRNAs in LBS single mutants (Fig S4B). By contrast, *BD mut* and *sygl-1(ø)* had virtually no cells with ≥5 mRNAs, but *BC mut* and *CD mut* both had a low percentage of cells with ≥5 mRNAs (Fig 5D). This cell-by-cell analysis reveals differences among the LBS double mutants that bulk measurements missed. Those differences are consistent with the conclusion that solo LBS C is less active than solo LBS B or LBS D.

### LBS number can function additively

Our initial analyses suggested that LBS number plays a critical role in shaping the *sygl-1* transcriptional gradient: three LBS generated a more abundant and extended gradient than two LBS, while two LBS generated a modest gradient in comparison and one LBS was close to null with no gradient (Figs 2-3). They also suggested that LBS number affects the extent of GSC maintenance. Both PZ size and GSC pool size were larger in wild-type than LBS single mutants, and both were gone in LBS double mutants (Fig 4). To further investigate the role of LBS number in shaping SYGL-1 output, we created animals heterozygous for distinct LBS regulatory regions (see Methods). To score molecular and biological outputs from one set of images, we placed all heterozygotes and all controls in an *lst-1(ø)* background.

We first compared *sygl-1* wild-type *(WT)* homozygotes to *WT/BCD mut* heterozygotes. *WT* carries six LBSs, three on each chromosome, whereas *WT/BCD mut* carries only three LBSs, three on the *WT* chromosome and none on the *BCD mut* chromosome. The three LBSs in *WT/BCD mut* produced 65% as much SYGL-1 as the six LBSs in *WT*, a bit more than half (Fig 6A). We next compared *WT* homozygotes with *B mut* homozygotes (four LBSs, two on each chromosome), and *B/BCD mut* heterozygotes (two LBSs, both on the same chromosome). The two LBSs in *B/BCD mut* produced about half as much as the four LBSs in *B mut*, and about a third as much as the six LBSs in *WT* (Fig. 6B). We were intrigued that SYGL-1 abundance in *B mut* was more than half of *WT* in *lst-1(ø),* but less than half of *WT* in *lst-1(+)* (compare Fig 6B to Fig 3B). This difference is explained by an effect of *lst-1(ø)* on SYGL-1 expression (Fig S7A-B). Regardless, when all mutants were scored in *lst-1(ø)* (Fig. 6B), LBS number or dose determines the relative SYGL-1 abundance, consistent with findings in *lst-1(+)* (Fig 2,3).

**Figure 6.**
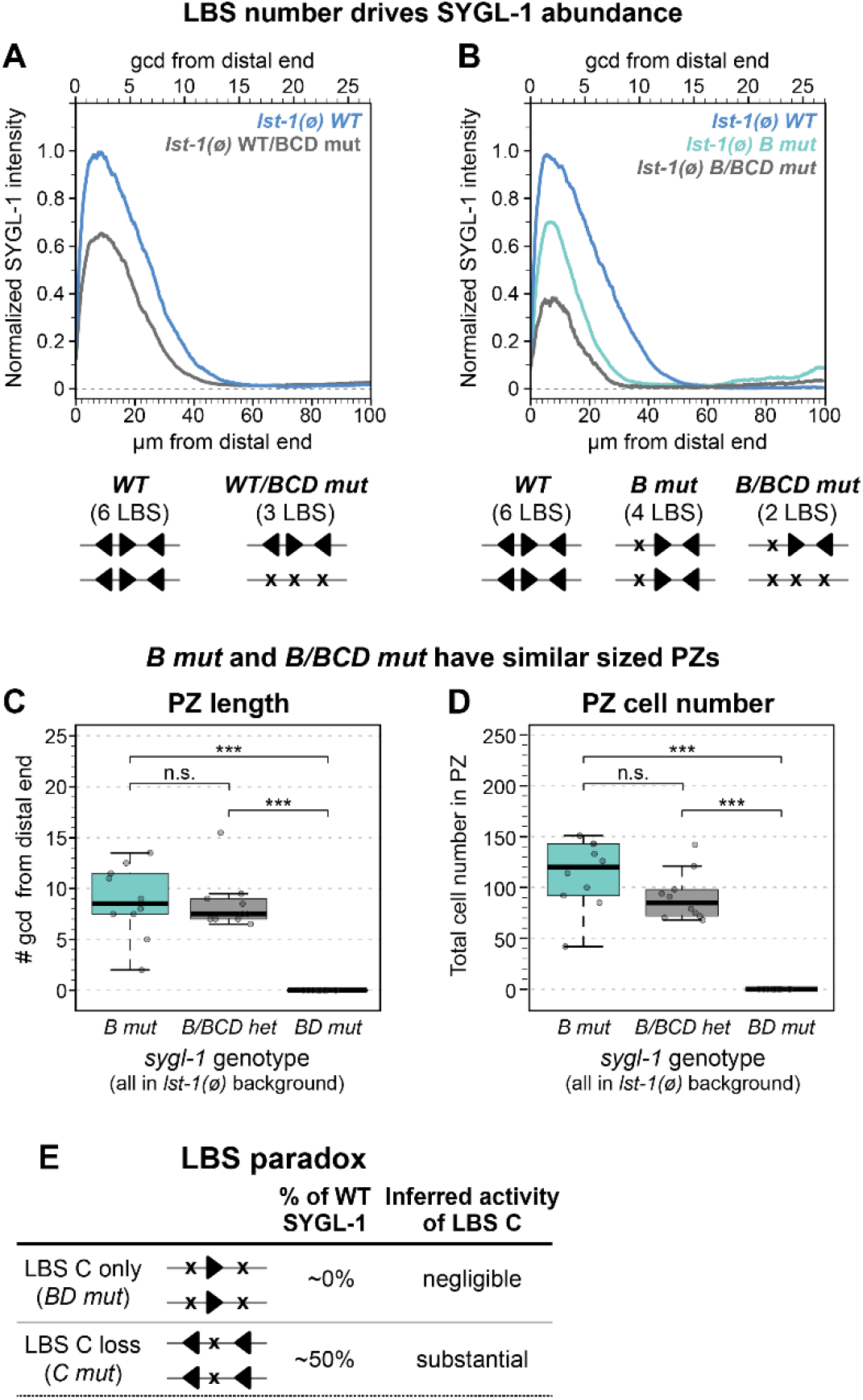
LBS number is a key factor in determining SYGL-1 abundance. **A-B.** Quantification of α-V5 immunofluorescence. Conventions as in Fig 3C, except only mean shown (Methods). See Methods for *sygl-1* heterozygote creation. **A.** 30 gonads per genotype scored in 2 experiments. **B.** Total gonads scored in 1-4 experiments: *wt*: 66; *B mut:* 73; *B/BCD mut:* 10; *BD mut:* 10. **C-D.** Boxplot conventions match Fig 2F. ***: P<0.01, n.s.: p≥0.01; Student’s two-tailed T-test (homoscedasticity assumed). 10 gonads per genotype scored from one experiment. **C.** Average # germ cell diameters (gcd) in PZ were counted by eye. p = 0.86 (*B mut* vs *B/BCD mut*). **D.** Total # cells in PZ counted using Imaris (see Methods). p = 0.11 (*B mut* vs *B/BCD mut*). **E.** Results in LBS single and double mutants are paradoxical; see text for explanation.

We next addressed the question of whether the SYGL-1 level generated in *B/BCD mut* was above or below the threshold for GSC maintenance. This question can only be addressed in a *lst-1(ø)* background where GSC maintenance depends solely on SYGL-1. All strains scored above for SYGL-1 abundance, including *B/BCD mut*, had a normal fertile germline with a PZ and GSCs maintained into adulthood. Thus, *B/BCD mut* makes SYGL-1 at or above the threshold required to maintain GSCs into adulthood. Unexpectedly, PZ sizes were similar in *B mut* with its four LBSs and *B/BCD mut* with its two LBSs (Fig 6C-D). The relationship between SYGL-1 abundance and PZ size must therefore be explored further, perhaps by creating additional LBS mutants that modulate SYGL-1 abundance more finely and analyzing in more depth the circuitry responsible for GSC maintenance.

### Neighboring LBS within a single cluster act synergistically

Comparison of results from LBS single mutants and LBS double mutants presented a paradox (Fig. 6E). A solo *sygl-1* LBS drove little or no expression in LBS double mutants and had little or no biological activity, but removal of any one LBS reduced expression substantially. For example, solo LBS C had no detectable activity in *BD mut* (Fig. 5B,D), but removal of LBS C in *C mut* had a substantial effect (Figs. 3, 4, S4B). How can an LBS impact *sygl-1* expression so much when mutated on its own, but have no activity when it is the only LBS retained in the cluster? We suggest that LBS synergy between neighboring LBS within a single cluster solves this paradox. In support of this idea, *B/BCD mut* heterozygotes possess only two LBSs, but they reside within the same cluster and make enough SYGL-1 to maintain adult GSCs (Fig. 6B-D). By contrast, *BD mut* homozygotes also possess only two LBSs, a solo LBS C in each cluster, but they are on different chromosomes and cannot maintain GSCs, even in larvae (Fig 5B, Fig 6C-D). Although SYGL-1 abundance could not be measured in *BD mut*, because it lacks GSCs and hence a germline tissue, we infer that SYGL-1 abundance in *BD mut* must be below the SYGL-1 threshold and is likely to be comparable to *sygl-1(ø)* (Fig. 5B). We conclude that neighboring LBS on the same chromosome act synergistically.

## Discussion

### Functional analysis of a homotypic CRE cluster in its natural developmental context

Clusters of closely spaced DNA *cis-*regulatory elements are common in animal genomes and point mutations in these elements are critical to evolution and disease (Crocker et al., 2016; Ezer et al., 2014; Gotea et al., 2010; Madani Tonekaboni et al., 2019; Melton et al., 2015; Payne and Wagner, 2015; Stern and Orgogozo, 2009). Traditionally, functional analyses of CREs and CRE clusters have relied on artificial assays, such as reporter transgenes and heterologous cells (e.g. Hardison and Taylor, 2012; Shlyueva et al., 2014), while analyses in their natural context remain largely unexplored. The *C. elegans sygl-1* LBS cluster has several advantages for *in vivo* analyses of CRE function. Rapid Cas9 gene editing in *C. elegans* and ability to freeze strains both facilitate the systematic generation of CRE mutants for comparison. In addition, the relative simplicity of *C. elegans* hits a sweet spot: mechanisms governing nematode transcription are metaozoan but with fewer layers of complexity than mammals. In this work, we investigate a homotypic cluster of three CREs in *sygl-1* regulatory DNA, each with the same conserved sequence. This cluster responds continuously to Notch signaling to drive *sygl-1* transcription and maintain germline stem cells throughout development. As a result, the *sygl-1* gene allows us to directly link LBS function to *sygl-1* transcription and stem cell maintenance. This simple Notch-dependent LBS cluster therefore provides a powerful model to understand how CREs and CRE clusters regulate development in time and space. Given the overwhelming importance of CRE clusters in developmental control and the recent advent of efficient gene editing, analyses of CRE function in a natural *in vivo* context will be an essential facet of future studies.

### LBS additivity and synergy

A cluster of three closely spaced Notch-dependent CREs, called LBS B – LBS D, drives *sygl-1* expression in the distal gonad. To explore activities of individual LBSs within the cluster, as well as LBS pairs, we generated a set of all possible LBS single, double and triple mutants. The three LBS single mutants reduced *sygl-1* expression dramatically with levels roughly equivalent to each other (Fig. 7A). Therefore, each LBS must be essential for generating the wild-type level of *sygl-1* expression, and the three must be roughly equivalent in activity. A corollary is that the pairs of wild-type LBSs remaining in each LBS single mutant have similar activities, despite differences in spacing and polarity (Fig. 7B). A previous study of a different *C. elegans* Notch-dependent *cis*-regulatory region, analyzed in embryos with a reporter transgene, also found no effect of LBS polarity on response strength (Neves et al., 2007). Thus, LBS activities within the *sygl-1* cluster can tolerate differences in orientation and spacing, though the limits of that tolerance have not been tested.

**Figure 7.**
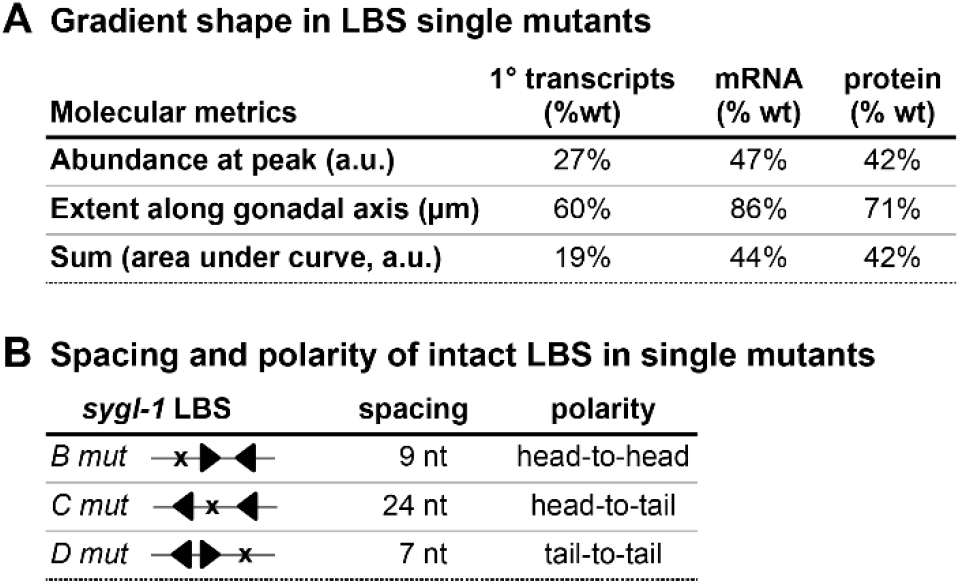
Summary of functional LBS analysis. **A.** Descriptors of gradient shape (as a percentage of wild-type): abundance at peak (0-5 μm from the distal end, varying units); extent (distance along gonadal axis in μm); sum (area under curve, in arbitrary units). *B mut, C mut*, and *D mut* values were averaged together into a single LBS single mut value (See Methods). 1°: primary (nascent) transcripts. **B.** Each pair of LBSs in single mutants drives similar synergy as any other pair, despite differences in spacing and polarity. Arrowhead conventions match Fig 1E.

Additional comparisons, including LBS homozygotes and heterozygotes, revealed that LBS activities are additive when on separate chromosomes (Fig 6A,B), but synergistic as neighbors within a single cluster (Fig 3F; Fig 6B-D). Notch-dependent CREs are also synergistic in flies and mammals (Arnett et al., 2010; Bailey and Posakony, 1995; Cave et al., 2005; Nam et al., 2007). Traditionally, CRE synergy is deduced when the transcriptional readout from neighboring CREs is substantially greater than the summed readout of separate CREs (Carey, 1991; Ptashne, 1988). To identify synergy between *C. elegans* LBSs, we compared *sygl-1* expression in animals carrying only two LBSs in *cis*, both on the same chromosome, to *sygl-1* expression in animals carrying only two LBSs in *trans*, one on each chromosome. Two LBSs in *cis* drove expression at a substantially higher level than two LBSs in *trans*, revealing synergy between neighboring LBSs. Indeed, the two LBSs in *cis* made enough SYGL-1 to maintain adult stem cells, whereas the two LBS in *trans* made too little to maintain stem cells. In flies and mammals, a head-to-head polarity of Notch-dependent CREs orients NICD proteins in neighboring transcription factor complexes to interact cooperatively at a molecular interface that has not been conserved in *C. elegans* (see Introduction). Because *C. elegans* LBS polarity is not critical (Neves et al., 2007; this work), it seems unlikely that *C. elegans* NICDs interact physically via a different interface. Therefore, we suspect LBS synergy employs a distinct mechanism.

One possibility for the molecular basis of LBS synergy in *C. elegans* is that neighboring LBSs enhance LAG-1/CSL occupancy at the cluster. Like many transcription factors, CSL proteins rapidly bind and release sites with dwell times on the order of one or a few seconds (Falo-Sanjuan et al., 2019; Giaimo et al., 2021; Gomez-Lamarca et al., 2018) and yet their transcriptional bursts last for much longer (10 – 70 minutes for *sygl-1)* (Falo-Sanjuan et al., 2019; Lammers et al., 2020; Lee et al., 2019). One can imagine that as LAG-1 releases from an LBS, the existence of a neighboring LBS increases its rebinding. This idea is consistent with the increased occupancy of CSL proteins observed upon Notch activation, especially in genes with clusters of binding sites (Castel et al., 2013; Housden et al., 2013; Wang et al., 2014). Another possibility is “assisted loading”, where binding of one LAG-1 protein to an LBS indirectly facilitates binding of another, for example by increasing chromatin accessibility and/or increasing the local concentration of other factors that aid LAG-1 recruitment and/or optimize its binding (Ezer et al., 2014; Falo-Sanjuan and Bray, 2020). Both possibilities are consistent with our finding that wild-type clusters with three LBSs drive more transcription than mutant clusters with two LBSs and dramatically more than clusters with only one LBS. They are also consistent with the finding that three neighboring LBSs on the wild-type chromosome in *WT/BCD mut* make as much SYGL-1 as two neighboring LBSs on each of the clusters in *B mut*, even though *WT/BCD mut* possesses one fewer LBS than *B mut* (three versus four) (Fig. 6A,B). Thus, both LBS number and neighborhood are important to activity level. An important question for the future is whether LBS mutants lead to differences in duration or frequency of *sygl-1* transcriptional bursts. Regardless, we suggest that the combination of LBS additivity and synergy reported here may apply to homotypic CRE clusters more broadly.

### SYGL-1 extent governs threshold for GSC self-renewal

An attractive concept is that the extent of SYGL-1 expression along the gonadal axis determines the size of the GSC pool (reviewed in Hubbard and Schedl, 2019). According to this model, GSCs self-renew in the presence of SYGL-1, but begin to differentiate upon its loss. This idea was proposed from the finding that expansion of the SYGL-1 gradient enlarges the GSC pool or can even form a tumor, depending on extent of the expansion (Shin et al., 2017). However, the effect of a smaller than normal SYGL-1 gradient had not been explored, because only *sygl-1* null mutants were available prior to this work. This work generates LBS mutants, which shrink the SYGL-1 gradient but do not abolish it. These mutants make enough SYGL-1 to maintain GSCs without its redundant counterpart LST-1, but they also reduce GSC pool size by about half. Therefore, an increase in SYGL-1 enlarges the pool (Shin et al., 2017), and a decrease in SYGL-1 shrinks it (this work). Together, these results show definitively that the extent of SYGL-1 expression along the gonadal axis determines GSC pool size.

GSCs are maintained where the SYGL-1 gradient is at its peak, and their daughters begin to differentiate where the gradient becomes too low. But how much SYGL-1 is sufficient to promote self-renewal and how low must SYGL-1 be to trigger differentiation? To gain insight, we measured SYGL-1 abundance in the absence of LST-1, the redundant counterpart of SYGL-1. When LST-1 is wild-type, stem cells can use either SYGL-1 or LST-1, effectively masking effects of varying SYGL-1 levels. However when LST-1 is removed, SYGL-1 alone is responsible for GSC maintenance. So the correlation between SYGL-1 abundance and stem cell maintenance was assayed in an *lst-1(ø)* background. We found that LBS single mutants and *WT/BCD mut* and *B/BCD mut* heterozygotes all made enough SYGL-1 to maintain adult stem cells and were thus above the threshold. By contrast, LBS double mutants made too little SYGL-1 to maintain adult stem cells and were thus below the threshold. Therefore, the SYGL-1 functional threshold is ≤40% of wild-type SYGL-1 abundance (Fig 6B). However, this number is based on measurements made in *lst-1(ø)* germlines and cannot be translated simply to *lst-1(+)* germlines (Fig S7).

### Future directions

This work manipulates CREs to tune the abundance of one crucial stem cell self-renewal regulator. A long term goal for comprehensive understanding of the decision to self-renew or differentiate will be to integrate cell-autonomous factors (e.g. abundance ratios between critical stem cell regulators like SYGL-1, LST-1 and their regulators) and non-cell-autonomous factors (e.g. mechanical force from tissue morphology, altered signaling in aged animals, the extracellular matrix, soma-germline communication) (Chacon-Martinez et al., 2018; Lin et al., 2020; Muncie and Weaver, 2018; Perez-Gonzalez et al., 2021; Starich and Greenstein, 2020). Untangling the relative contributions of these factors is a next challenge to improve predictive models and stem cell therapies. This work modulates SYGL-1 abundance using LBS manipulations and lays a foundation in a tractable model organism to test a variety of genetic and environmental contexts and begin to tease apart these critical ratios and relationships.

## Acknowledgements

The authors thank David Wassarman and members of the Kimble and Wickens labs for helpful discussions throughout the course of this work. We thank Jane Selegue, Peggy Kroll-Conner, and Hannah Moulton for assistance with experiments or strain building. We thank Ahlan Ferdous, Sarah Crittenden, Melissa Harrison and David Wassarman for critical comments on the manuscript. Thanks to Sam Engle and the UW-Madison CALS Statistical Consulting Lab for discussions on statistics. TRL was supported by the NSF; this material is based upon work supported by the National Science Foundation Graduate Research Fellowship under Grant No. DGE-1747503. Any opinions, findings, and conclusions or recommendations expressed in this material are those of the authors and do not reflect the views of the National Science Foundation. JK was an Investigator of the Howard Hughes Medical Institute and is now supported by NIH R01 GM134119.

## Author contributions

TRL conceived and performed experiments, analyzed data, and wrote the paper. MX and CC performed experiments and analyzed data. CHL analyzed data. JK conceived experiments and wrote the paper.

## Methods

### *C. elegans s*trains and nomenclature

See Table S1 for a list of genotypes and the figures each strain was used in. “*ø*” refers to null mutants that have a CRISPR deletion of the coding region; *“+”* refers to a wild-type copy of the gene. For simplicity, several different genotypes may be referred to using the same in-text abbreviation (e.g. “*D mut”* can refer to *lst-1(ø) sygl-1::V5(D mut), lst-1(ø) sygl-1(D mut), sygl-1(D mut),* or *lst-1(ø) sygl-1::V5(D mut); emb-30(tn377)).* All strains were maintained at 20°C unless noted, using standard culture techniques (Brenner, 1974). The balancer used for *LG I* was *hT2[qIs48]* (Siegfried and Kimble, 2002).

### CRISPR/Cas9 gene editing

No transgenes were used; all genetic edits were done by CRISPR/Cas9 genome editing at the endogenous loci. Recombinant Cas9 protein (Paix et al., 2015), single-stranded DNA oligo repair templates complementary to the nontarget strand (Richardson et al., 2016), and custom crRNAs and tracrRNA (Integrated DNA Technologies) were injected into the *C. elegans* germline. A co-conversion approach was used (Arribere et al., 2014): single-stranded DNA repair templates and crRNAs were designed against both the *sygl-1* mutation of interest and a *dpy-10* co-injection marker. Edits of the *dpy-10* co-injection marker create visible phenotypes, decreasing the number of animals that must be screened for the desired *sygl-1* edit by PCR. The *dpy-10* repair and crRNA sequences are from the Fire Lab (repair: 5’ CACTTGAACTTCAATACGGCAAGATGAGAATGACTGGAAACCGTACCGCATGCGGTGCCTATGGTAGCGGAGCTTCACATGGCTTCAGACCAACAGCCTAT 3’; crRNA: 5’ GCTACCATAGGCACCACGAG 3’). Final concentrations in the injection mix are as follows: *dpy-10* repair 1.34 μm; *dpy-10* crRNA 4 μm; gene-specific repair 4 μm; gene-specific crRNA 9.6 μm; tracrRNA 13.6 μm; recombinant Cas9 protein 25 μm).

### Scarless CRISPR/Cas9 genome editing

To create a canonical LBS A motif (Fig S1C-D), we followed a two-step CRISPR-Cas9 editing protocol which in the first step creates the point mutation and replaces a stretch of 23 nt in the *sygl-1* 5’ flanking sequence with *dpy-10* protospacer sequence; the second step removes the *dpy-10* sequence to create a scarless edit (El Mouridi et al., 2017). After completing the first step of this protocol, we found that mutating LBS A to a canonical LBS produced SYGL-1 at levels similar to wild-type and abandoned the second step.

### Design of LBS mutation

Both the 5’ TGACGTCA 3’ and 5’ GGATCCAA 3’ LBS mutations were designed to mimic LBS mutations from the literature where loss of function had been demonstrated (Choi et al., 2013; Christensen et al., 1996; Kershner et al., 2014; Neves et al., 2007; Yoo et al., 2004). Mutating the endogenous LBSs presented a genotyping challenge in that the nucleotides in the LBS mutation are the only change from wild-type. Thus, we mutated five nucleotides and designed mutant sequences that add a restriction enzyme cleavage site *(AatII* for 5’ TGACGTCA 3’ and *BamHI* for 5’ GGATCCAA 3’). Additionally, LBS mutation disrupts the PAM sequence. We alternatively found that designing PCR primers whose 3’ ends overlapped with the mutated five nucleotides successfully generated mutant-specific or wild-type-specific PCR product.

### *sygl-1* gene diagrams

In Figure 2A, we show a predicted transcription start site (TSS); this TSS was drawn using the Wormbase version WS280 annotation and RNA seq data.

Gene diagrams of *sygl-1* are all drawn to scale except that smFISH probes in Fig 2A are drawn larger than to scale for visibility; the center of each line indicates the 5’ end of each 20nt-long smFISH probe.

### Immunostaining

#### General staining protocol

Immunostaining was performed as described by the solution extrusion method in Crittenden et al., 2017 with minor modifications. Specific protocol adaptations are listed in sections below. Unless otherwise noted, animals were dissected at L4 + 24 hours at 20°C in a glass petri dish with ~10 mL of PBSTw+0.25 mM levamisole. Samples were moved to a 1.5 mL Eppi tube, fixed in paraformaldehyde (Thermo 28908), washed in 1mL PBSTw, then permeabilized and washed briefly three times in PBSTw before blocking in PBSTw+0.5% BSA (PBSB) for at least 30 minutes at room temperature. Samples were incubated in the primary antibody solution at 4°C overnight, then washed 3-4 times in 1 mL PBSTw before adding 100-200 μL of secondary antibody solution. Samples were incubated in the dark at room temperature for 1 hour. Another series of 3-4 PBSTw washes was repeated, shielding samples from light, then samples were mounted in ProLong Gold antifade reagent (Fisher P36930) on glass slides (FisherFinest Premium 12-544-1) with 22×22 mm coverslips (Azer Scientific ES0107052) and cured in a dark drawer at room temperature prior to imaging (cure times overnight to several days).

#### V5 epitope tag staining

Fixation: 3% paraformaldehyde for 20 minutes; permeabilization: 0.5% triton in PBS for 5 min; Mouse anti-V5 antibody (SV5-Pk1, Bio-Rad, MCA1360, 1 mg/mL, Dunn et al. *J Immunol methods* 1999) was diluted 1:1000 in PBSB; Secondary antibody solution: 0.1 μg/mL DAPI + Donkey anti-Mouse Alexa 555 (1:1000, Thermo Fisher Scientific A31570, lot 1117032); antibody solutions were removed with two quick (10 inversions of tube) and two long (10-15 min on rocker) washes. Mounted in 12 μL ProLong Gold Antifade (Thermo Fisher Scientific P36930 or P10144). Note: the α-V5 antibody non-specifically stains the distal tip cell body.

#### GLD-1 and PH3 staining

The same staining protocol was used for GLD-1 staining (Fig 4E) and *emb-30* assay (Fig 4F-G) except that in the *emb-30* assay animals were maintained at different temperatures and staged differently (see *emb-30* assay). Note: Fig 4B cartoon GLD-1 staining is a representation of wild-type animals, not the *lst-1(ø)* control animals. This work is consistent with previously published GLD-1 staining in *lst-1* null animals; GLD-1 is brighter in the distalmost region and shifted distally compared to wild-type (Brenner and Schedl et al. 2016).

Fixation: 4% paraformaldehyde for 10 minutes; permeabilization: 0.5% triton in PBSTw+0.5%BSA (PBSB) for 10 minutes; Rabbit anti-GLD-1 (Schedl Lab, Jones et al., 1996) was diluted 1:200 in PBSB; Mouse anti-Phospho-Histone H3 (Ser10) (Cell signaling 9706L, lot 10) was diluted 1:200 in PBSB. Secondary antibody solution: 0.1 μg/mL DAPI and Donkey anti-Rabbit Alexa 647 (Invitrogen Molecular Probes A31571 lot 1252811), Donkey anti-Mouse Alexa 488 (Invitrogen Molecular Probes A21202 lot 1226927). Antibody solutions were removed with two quick (10 inversions of tube) and one long (15 min) wash. Mounted in 18 μL ProLong Glass (Fisher, P36984)

#### Immunostaining for progenitor zone analysis

For progenitor zone size counts, a similar protocol as above was used except animals were fixed for 15 min in 2% paraformaldehyde, permeabilized 15 minutes in 0.5% triton, blocked for 30+ minutes, and stained with mouse anti-DAO-5 1:100 (DSHB, Hadwiger et al., 2010). Some progenitor zone length counts used a shortened method of the above protocol: fixation for 10 minutes in 4% paraformaldehyde + PBSTw, followed by one quick PBSTw wash, 5 minute permeabilization in 0.1% triton+PBS, then one wash in PBSTw and mounted in Vectashield + DAPI (Vector Labs, H-1200).

#### Image Acquisition

Images were captured on a Leica SP8 confocal microscope. Gonads were imaged from top to bottom with a z slice depth of 0.5 μm. Channels were acquired sequentially (between stacks). α-V5 signal was excited at 561 nm (0.5%, HeNe) and signal was acquired from 564-615 nm (gain 70) with 16 line averages and 2 frame accumulations using a 63x objective at 150% zoom and an 8000 Hz scan head; α-GLD-1 signal was excited at 633 nm (0.2%) and acquired from 564-614 nm (gain 70, 2 line averages) using a 40x objective and 125% zoom or a 63x objective and 100% zoom and a 400 Hz scan head; α-PH3 signal was excited with 561 nm (0.15%, Argon at 25% 24W) and collected from 642-693 nm (gain 80, 1 line average) using a 63x objective and 100% zoom and a 400 Hz scan head; DAPI was excited at 405 nm (0.8-1.2%, UV) and signal acquired from 412-508 nm (gain 500-700).

### Fluorescence quantitation

FIJI (<u>F</u>IJI <u>I</u>s <u>J</u>ust <u>I</u>mageJ) was used to quantitate fluorescence pixel intensities. Workflow is as described in Haupt et al. 2019 and Brenner et al. 2016. A FIJI macro was written to automatically create sum Z projections and save the output as TIFFs. A 50-pixel wide freehand line was drawn from the DTC down the germline image. FIJI Plot Profile tool was used to copy pixel intensity data and paste into Microsoft Excel. *Background subtraction*

In Microsoft Excel, an average pixel intensity for each genotype was created by averaging values from individual germlines at each x coordinate. Then, the average pixel intensity per genotype was background subtracted using the average pixel intensity from the negative control (untagged) sample; the exception is Figure 1, where the untagged JK5622 *sygl-1(q828)* control values are not subtracted in order to display them in comparison to *BCD mut* values. For all other experiments, the untagged control was N2 (wild-type).

#### Normalization of background-subtracted pixel intensities

After pixel intensities were processed as above, intensities were normalized in one of two ways. For Figures 1 and 3, the values at each x coordinate were divided by the maximum intensity from the positive control to transform intensities into a percent of control.

For the GLD-1 data in Fig 4E, processed pixel intensities were normalized against an internal control: JK4864 (Table S1) worms expressing a GFP reporter in somatic cells were grown on the same plates and dissected in the same dish as the test worms. The JK4864 control worms have normal GLD-1 expression and the peak value in the JK4864 data was set to 1.0 for each slide.

#### SEM shading on plots

The SEM values were calculated on background-subtracted pixel intensity values; values from different experiments were averaged together and then normalized. Standard error was calculated using the number of experiments as n. Thus, the SEM shading reflects the spread of pixel intensity values on different slides imaged on different days.

SEM shading was used on immunostaining data for some figures but not others. For SYGL-1 abundance figures lacking SEM shading, we normalized background-subtracted values to their respective controls (stained in parallel) before averaging together values from different replicates, which makes mean values robust against day-to-day variation in pixel intensity.

#### Total protein (area under the curve)

To calculate total protein quantitation (Fig 3D), arrays X (distance values in μm) and Y (normalized SYGL-1 intensity values) were imported into MATLAB version R2015a and the command trapz(X,Y) was used to calculate area under the curve.

#### Measurements along gonadal axis

Microns were converted to germ cell diameters (gcd) using a conversion factor of 3.7 μm/gcd, which was calculated by manually measuring cell diameters in FIJI (10 cells were measured for 1 randomly chosen image from each experiment, using both protein staining and smFISH experiments).

### Single molecule RNA FISH (smFISH)

The same *sygl-1* exon and intron probes were used as described in Lee et al, 2016. See also a published smFISH protocol (Lee et al., 2017). Briefly, mid-L4 stage animals were grown on OP50 at 20°C for 24 hours, then dissected as described in Immunostaining. Animals were fixed in 3.7% formaldehyde (37% formaldehyde Amresco, 0493-500ML) for 15-25 minutes and permeabilized in RNAse-free PBS (Fisher BP24384) + 0.1% triton for 10-12 minutes. After an overnight incubation in 70% ethanol (diluted with DEP-C-treated H2O (Ambion AM9922), samples were equilibrated in smFISH wash buffer for 15-20 minutes, then incubated in hybridization buffer plus smFISH probe at 37°C for 46-48 hours. Lyophilized smFISH probes were resuspended in RNAse-free TE buffer (10 mM Tris-HCl, 1 mM EDTA, pH 8.0) to make a 250 μM stock solution. The *sygl-1* exon-specific probe set includes 31 unique oligonucleotides labeled with CAL Fluor Red 610 and was used at a final concentration of 0.25 μM. The *sygl-1* intron-specific probe set includes 48 unique oligonucleotides tagged with Quasar 570 and was used at a final concentration of 0.50 μM. Samples hybridized at 37°C overnight, then were washed with smFISH wash buffer and 1μg/uL DAPI at 37°C for 40-50 minutes. Finally, samples were resuspended in 12 μL Prolong Gold antifade reagent (Fisher P36930), mounted on glass slides, and cured unsealed in a dark drawer for at least 24 hours to several days before imaging.

#### Image acquisition settings

Images were captured on a Leica SP8 confocal microscope with the same hardware and Leica software as described in Lee et al., 2016. Gonads were imaged from top to bottom with a z slice depth of 0.3 μm, with a 63x objective at 300% zoom. Channels were acquired sequentially in the following order: *sygl-1* intron Q570 probe was excited at 561 nm (3%, HeNe) and signal was acquired from 564-588 nm (gain 40); *sygl-1* exon C610 probe was excited at 594 nm (3%, DPSS) and signal acquired from 600-680 nm (gain 40); DAPI was excited at 405 nm (0.8-1.2%, UV) and signal acquired from 412-508 nm (gain 500-700). Six of nine experiments were imaged between frames with a 400 Hz scan head and line average of 6 for both RNA channels or 3 for DAPI; three of nine experiments were imaged between stacks with an 8000 Hz scan head and line average of 16 for intron, 32 for exon, and 8 for DAPI with 2 frame accumulations on each channel.

#### Representative images

Image contrast was adjusted in FIJI equivalently for all main images and for all insets. Contrasts for intron channel differ between main images and insets to highlight ATS number in main images and ATS intensity in insets. Exon channel is pseudo-colored with FIJI default Fire LUT to show cytoplasmic mRNA without saturating ATS. All gonad images were taken from the same experiment.

### MATLAB analysis of smFISH data

#### Removal of data points

Some images in a few experiments displayed bright, nonspecific signal outside the gonad tissue that interfered with detection. Images where the nonspecific signal was greater than 8 um^2^ were removed (1 *wt*, 10 *C mut*, 6 *D mut*, 3 *CD mut*, 1 *sygl-1(ø)).* Additionally, a small number of images threw errors in MATLAB related to reading the image files (e.g. nuclei imaged too close to edge of frame) and were also removed (5 *C mut*, 5 *BD mut*, 3 *CD mut*, 1 *sygl-1(ø)).* Total number of images quantified are reported in Fig S2E.

#### MATLAB code

For MATLAB code analysis, refer to Lee et al., 2016, Crittenden et al., 2019. Code modifications described in Crittenden et al., 2019 were also applied to this work. Threshold values were set as 1.0 thresForIntron and 0.55 thresForExon for all images. To define cell boundaries for #mRNA/cell calculations, a Voronoi with 3 μm limit was used (the same as in Crittenden et al. 2019 and Lee et al. 2016; does not include center of rachis).

#### Cytoplasmic spots versus mRNA

The dim cytoplasmic spots range slightly in intensity and size. For *wt* and each LBS mutant, ~60% of cytoplasmic spots have a dim intensity distribution with a low coefficient of variation, consistent with a single mRNA (Fig S3A) (see also Lee et al., 2016). Approximately 1/3 of the cytoplasmic spots in each genotype are a bit larger and their intensity values are approximately twice that of the single mRNA population, consistent with two mRNAs. Rare spots have ≥3 mRNA per cytoplasmic spot. Where possible, number of mRNA are reported rather than the number of cytoplasmic spots, but the #mRNA/cell values represented in Fig 3B, 3E, 5D and S4B are the number of cytoplasmic spots.

#### Background level and extent of sygl-1 mRNA gradient

The background level in Figures 3B and 3E is 0.956 cytoplasmic spots (or ~1.25 mRNA). This background was determined by calculating one standard deviation above the mean number of cytoplasmic spots detected in the 45-50 μm region of *sygl-1(ø)* control gonads. Local minimums indicate the end of the Notch-dependent *sygl-1* gradient because the slight increase in RNA in the 40-60 μm region is consistent with the rise in proximal Notch-independent *sygl-1* expression.

#### Normalization of ATS pixel intensities

To control for image-to-image intensity changes, raw exon channel ATS values from each gonad are divided by the mean intensity for the raw single mRNA population (the population of cytoplasmic spots estimated to be one mRNA) within the same gonad. Then, the mean of all the single mRNAs is transformed to 10. Thus, a normalized ATS intensity of 50 was five times brighter than the mRNA within the same image. Normalized ATS intensities were divided by 10 to estimate the number of nascent transcripts per ATS (Fig 2F).

#### Number of ATS per nucleus pie charts (Fig. 2D)

The percentages of nuclei containing x number of ATS were calculated individually for each experiment. The percentages in the pie charts are the average percent of all the experiments in the datasets. For example, the LBS double mutant dataset contains 94 nuclei and only one of those nuclei contains 3 ATS. That 3-ATS nucleus came from one experiment with 21 total nuclei; that experiment’s 3-ATS percentage is 4.8%. The other 7 experiments that contained any ATS-positive nuclei had 0% nuclei with 3 ATS and thus the overall average is 0.6%.

#### Summary of sygl-1 mRNA data (Fig 3D)

The total number of single mRNAs per experiment was divided by the number of gonads in that experiment; standard deviation was calculated between experimental averages. The peak number of mRNA is the maximum number of cytoplasmic spots detected (bar height in 0-5 μm bin of ave #mRNA/cell plots) as a ratio to WT; standard deviations were calculated between the average values for each independent experiment. Total # mRNA: Ave # mRNA as described above as a ratio to WT.

### Germline function

#### Percent fertile

To score percent fertile, F1 homozygous L4s were picked from balanced LBS single mutant stocks onto a fresh plate. 24 and 48 hours later, one L4+24 or L4+48 adult was singled onto a plate to lay embryos for 1.5-3.5 hours. Percent fertility of F2 homozygotes was scored (gravid or not fertile) at low magnification on a Zeiss Discovery.V12 microscope four days after eggs laid. Progeny from a total of eight different adults per genotype were scored. LBS double mutant animals were maintained for many generations and a gravid homozygous animal was never seen. Additionally, *lst-1(ø) sygl-1(LBS double mut)* homozygous L4s were raised at 15° C, 20° C, and 25°C and failed to produce embryos at all temperatures.

#### Percent Glp

To score percent Glp (germline proliferation abnormal), L4 + 24hr LBS double mutant adults were assessed at high magnification on a Zeiss Axio Imager.D1 microscope by DIC. Sperm were observed at the distal tip of all LBS double mutant germlines at 20° C. LBS single mutants are 0% Glp because 100% of germlines have a progenitor zone.

### Progenitor zone size

Progenitor zone size was assessed in animals grown at 20°C for 24 hours past mid-L4. Gonads were extruded, fixed, and stained with DAPI or DAPI and α-DAO-5 antibodies (see Immunostaining).

#### Progenitor zone length

PZ lengths were determined as follows: first, the DTC was located, then the number of cell rows between the DTC and the first crescent were counted. Both sides of the germline were counted and averaged together for one PZ length per germline. Note that because a curved metaphase plate can look similar to a crescent cell, the definition of “first crescent” cell included a requirement for multiple nearby crescent cells. For more information on identifying and scoring PZ, see (Crittenden et al., 2017). Gonads were visualized on a computer monitor for ease of manual counting; μManager software (a plugin for ImageJ) was used to connect the camera and computer (Edelstein et al., 2010).

#### Number of cells in progenitor zone

A protocol using Imaris software was modified from (Braude et al., 2015; Gopal et al., 2017; Sadeghi et al., 2014). A surface was drawn to mask the PZ region using the drawing contour mode with 1.7 um vertex spacing. The PZ boundary along the gonadal axis was chosen by eye on a middle slice where the first crescent cells could be seen, then the outline was copy/pasted to the first and last slices to create a 3D surface. The surface was used to mask the DAO-5 channel. A spots function with an estimated XY diameter of 2.0 um was created in the masked DAO-5 channel. The number of spots was filtered with a quality filter and the threshold was set by moving the slider until no spots were detected outside the germline. Detection was briefly visually checked and the number of spots detected was recorded as the number of cells in the PZ. We also used Imaris to count the number of cells in the PZ for some experiments that were DAPI stained but not DAO-5 stained (Fig S5A-B, Fig 7E). The protocol is the same except we used a 2.7 estimated XY diameter on a masked DAPI channel and spent more time per gonad checking and editing detected spots.

Imaris version 9.3.1 was used on a Dell Precision 5820 with 64-bit Windows 10 Education operating system, an Intel® Xeon® W-1245 CPU @3.70GHz processor, and 128 GB of RAM. Leica .lif images were converted to .ims images using Imaris FileConverter v9.5.0.

### Sperm counts

Mid-L4 larvae were harvested and whole-mount stained with the reduction/oxidation method (Finney and Ruvkun, 1990). Samples were fixed in Ruvkun fixation buffer with 1% paraformaldehyde for 30 minutes, followed by disulfide linkage reduction and incubation in blocking solution [1X PBS with 1% bovine serum albumin, 0.5% Triton X-100, 1mM EDTA] for 40 minutes. Samples were then incubated overnight at 4°C with rabbit α-SP56 (1:200, Ward et al. 1986) in blocking solution. Secondary antibody Alexa 555 Goat anti-rabbit (1:1000, Invitrogen Molecular Probes A21429, lot 1246456) was diluted in blocking solution and incubated with samples for at least 2 hours. Samples were washed with blocking solution and 1ng/μL DAPI for 15 minutes, then mounted in Vectashield (#H1000; Vector Laboratories, Burlingame, CA) for visualization with fluorescent compound microscopy.

#### Calculation of germ cell number

Germlines lacking both *lst-1* and *sygl-1* begin precociously differentiating in L1, two full larval stages earlier than L4, and have completed both spermatogenic meiotic divisions by L3, which creates 16-32 sperm from a total of 4-8 germ cells (Kershner et al., 2014). The distalmost cells were scored as undifferentiated by DAPI morphology or as differentiating (either sperm by DAPI morphology and SP56+ staining or meiotic by DAPI morphology). Estimates of the number of germ cells were extrapolated as follows: ¼ GSC per sperm, ½ GSC per secondary spermatocyte, and 1 GSC per primary spermatocyte or meiotic cell. Sperm and spermatocytes were counted in one gonad arm in JK5911 and JK6180 and multiplied by two. Sperm were counted in both gonad arms in JK6165 and JK6401.

### Creating *sygl-1* heterozygotes

LBS mutant/null heterozygotes were created by mating males of an LBS mutant with *sygl-1* null hermaphrodites. Heterozygote cross progeny were assayed by staining for α-V5: SYGL-1 protein is quicker to image and quantify than *sygl-1* mRNA, and LBS effects on SYGL-1 protein abundance were similar to those on *sygl-1* RNA abundance. *BCD mut* was chosen as the *sygl-1* null allele rather than *sygl-1(ø)* (Fig S6A-C). All heterozygotes were created in an *lst-1(ø)* background: SYGL-1 minimum threshold is best defined in *lst-1(ø);* GSC maintenance and SYGL-1 abundance could be tested in parallel; absence of *lst-1* allowed reliable detection of cross progeny (see below).

#### Reliably distinguishing cross progeny

Hermaphrodite parents in the heterozygote crosses throw 100% sterile progeny. Therefore, any progeny that maintain GSCs as adults must be heterozygote cross progeny. Only a small percent of progeny on successful mating plates were sterile, which is consistent with the idea that all sterile progeny are self-fertilized progeny from mother hermaphrodites. However, we decided to additionally rule out the possibility that some cross progeny are sterile by using a recessive *rol-6(e187)* allele: only self-fertilized progeny display a Roller phenotype. Thus, heterozygous cross progeny were additionally distinguished as wild-type movers. We repeated a heterozygous cross three times and found that 100% of wild-type mover progeny maintained GSCs; we concluded that it is unlikely the heterozygous cross progeny population includes any sterile animals. After this proof of principle, we assumed that sterile Glp animals were self progeny for all other heterozygous crosses due to technical practicalities.

#### Strains mated for crosses (see also Table S1)

In Figure 6B-D, *B/BCD mut* are progeny from JK6517 males mated with JK6600 hermaphrodites. In Figure 6A, *WT/BCD mut* are progeny from JK6431 males mated with JK6600 hermaphrodites. In Fig. S6B, *WT/ø* are progeny from JK6431 males mated with JK6401 hermaphrodites. In Figure 6A-D and Fig S6, control animals were also progeny from a male x hermaphrodite cross (e.g. *B mut* in Fig. 6B are progeny from JK6517 males mated with JK6517 hermaphrodites).

#### *emb-30* assay for number of germline stem cells

The *emb-30* assay to estimate the naïve GSC pool size was done as previously described (Cinquin et al. 2010). Animals with *emb-30(ts)* alleles were grown at 15°C permissive temperature until 36 hours past L4, then shifted to 25°C restrictive temperature for 12.5 hours. Temperature shifts were done in a programmable incubator (Echotherm IN35, Torrey Pines Scientific). Gonads were dissected, fixed, and stained for GLD-1, PH3, and DAPI (see Immunostaining). The number of naïve GSCs was estimated by using the multi-point tool in FIJI (Schindelin et al., 2012) to count the number of nuclei distal to the GLD-1 staining boundary that were positive for PH3 signal and/or mitotic by DAPI.

#### Selection of data for analysis

Six of 17 control *lst-1(ø) sygl-1::V5(D mut); emb-30* gonads, kept at permissive temperature, were meiotic. None of the animals in the *lst-1(ø) sygl-1::V5(wt); emb-30* controls (permissive and restrictive) were meiotic, so we interpreted this as a consequence of the mutant *emb-30* background and discarded all meiotic gonads (5/21 for *D mut*, 0/28 for *wt*) from analysis. As established in previously published *emb-30* counts (Crittenden et al., 2019), temperature-shifted gonads where most germ cell nuclei were fragmented (4/21 for D mut, 2/28 for WT) were also excluded due to unreliable counting.

### Descriptors of gradient shape

Values for Figure 7A summary were calculated by averaging together the individual *B mut, C mut*, and *D mut* values. We did so because when we consider all factors (e.g. directness of comparison in experimental design, consistency across multiple assays), our clearest conclusions tended to focus on similarities rather than differences between individual LBS single mutants. When we took the same factors into account for LBS double mutants, we found that strong conclusions extended to a clear distinction between *BD mut* and the other two LBS double mutants.

#### Calculation of a summary LBS single mutant gradient

Primary transcript values are as follows: abundance at peak values from % cells with *sygl-1* ATS Fig 2C graphs (LBS mut: 14.2 ± 5%; WT: 51.8 ± 8%); extent values from Fig 2C graph vertical dashed lines (LBS mut: 15 μm; WT: 25 μm); sum values are an estimate of the total number of primary transcripts per gonad using numbers from Fig S2E (LBS mut: 5.8 ± 4.1 ATS/gonad * 4.5 ± 0.6 tx/ATS = 17 tx/gonad; WT: 24 ± 8.5 ATS/gonad * 5.6 ± 0.5 tx/ATS = 134 tx/gonad). The mRNA values are: abundance from Fig 3A,C (LBS mut: 29 ± 3 mRNA at peak; WT: 62 ± 5 mRNA at peak); extent from Fig 3A (LBS mut: 30 μm; WT: 35 μm); sum from Fig 3C (LBS mut: 2616 ± 864 mRNA; WT: 6274 ± 1478 mRNA). The protein values are: abundance from Fig 3D (LBS mut: 42 ± 13% of WT peak); extent from Fig 3B (LBS mut: 43 μm; WT: 52 μm); sum from Fig 3D (LBS mut: 11557 ± 3678 a.u.; WT: 2781 ± 5739).

### Statistics

#### Box plots

Box plots were generated with the BoxPlotR web tool (Spitzer et al., 2014). BoxPlotR conventions: box limit: 25^th^ and 75th percentiles; whiskers extend 1.5 times the interquartile range from the 25^th^ and 75^th^ percentiles.

#### Lme4 package in R

Student T-Test (T.TEST function in Excel) was used whenever two samples could be compared. For multiple pairwise comparisons, we chose a linear mixed effects model. Observations were fitted to a linear mixed effects model (lmer) using lme4 package. R version 4.0.5 (2021-03-31) -- “Shake and Throw” and RStudio version 1.4.1106 “Tiger Daylily” were used with Windows 10. The genotype was the fixed effect and the experiment block was the random effect. The emmeans package was used to make pairwise comparisons between genotypes for a given bin.

#### Standard Error bars

We used the number of gonads to calculate SEM bars for *sygl-1* mRNA data because there are multiple mRNA/cell measurements from a single gonad. In all other cases, we used the number of experiments to calculate SEM bars because there was only one measurement per gonad.

