## Supplemental for "Notch-dependent DNA *cis*-regulatory elements and their dose-dependent control of *C. elegans* stem cell self-renewal"

Table S1: Strains used

| **Strain** | **Source** | **Description** | **Used in Figures** |
|---|---|---|---|
| N2 bristol | Brenner, 1974 | Wild-type | 2B-F; 3A-F; 5A,C; S2-S4 |
| JK4864 *qIs147* | | *Psur5::GFP* | 4E |
| JK5622 *q828 I* | Shin, 2017 | *sygl-1(ø)* | 1F-G; 2B,C,E; 3A,C,E; 5A,C; S1B; S2E; S3A; S4A,C; S6B-C |
| JK5773 *q936 I* | this work | *sygl-1(LBS C mut)* | 2C-F; 3A,C; S2B,D,E; S3A; S4A,C |
| JK5796 *q869 I/hT2[qIs48](I;III)* | Shin, 2017 | *lst-1(ø)* | 4A,C,D; 6A-C; S5; S6B-C |
| JK5812 *q942 I* | this work | *sygl-1(LBS CD mut)* | 2C-F; 3E; 5A,C; S2C-E; S3A; S4A |
| JK5813 *q943 I* | this work | *sygl-1(LBS D mut)* | 2B-F; 3A,C; S2B,D,E; S3A; S4A,C; S5C,D |
| JK5911 *q869 q942 I/hT2[qIs48](I;III)* | this work | *lst-1(ø) sygl-1(LBS CD mut)* | 4A; 5B |
| JK5912 *q956 I* | this work | *sygl-1(LBS B mut)* | 2C-F; 3A,C; S2B,D,E; S3A; S4A,C |
| JK6002 *q1015 I* | Shin, 2017 | *sygl-1::1xV5* | 1F-G; 3B,D,F; 6A; S1B,D; S3B |
| JK6020 *q869 q943 I/hT2[qIs48](I;III)* | this work | *lst-1(ø) sygl-1(LBS D mut)* | 4A,C,D; S5B,E-F |
| JK6065 *q1039 I* | this work | *sygl-1(LBS BD mut)* | 2C-F; 3E; 5A,C; S2C-E; S3A; S4A |
| JK6111 *q1054 I* | this work | *sygl-1::1xV5(LBS D mut)* | 3B-C; 6A; S3B |
| JK6161 *q1101 I* | this work | *sygl-1(LBS BC mut)* | 2B-F; 3E; 6A,C; S2C-E; S3A; S4A |
| JK6165 *q869 q1039 I/hT2[qIs48](I;III)* | this work | *lst-1(ø) sygl-1(LBS BD mut)* | 4A; 5B |
| JK6180 *q869 q1101 I/hT2[qIs48](I;III)* | this work | *lst-1(ø) sygl-1(LBS BC mut)* | 4A; 5B |
| JK6219 *q869 q936 I/hT2[qIs48](I;III)* | this work | *lst-1(ø) sygl-1(LBS C mut)* | 4A,C,D |
| JK6288 *q1135 I* | this work | *sygl-1::1xV5(LBS BC mut)* | 3F; S3C |
| JK6289 *q1136 I* | this work | *sygl-1::1xV5(LBS BD mut)* | 3F; S3C |
| JK6387 *q1163 I* | this work | *sygl-1::1xV5(LBS A mut)* | 1F-G; S1B |
| JK6388 *q1165 I* | this work | *sygl-1::1xV5(LBS C mut)* | 3B,D; S3B |
| JK6389 *q1167 I* | this work | *sygl-1::1xV5(LBS BCD mut)* | 1F-G; S1B |
| JK6390 *q869 q956 I/hT2[qIs48](I;III)* | this work | *lst-1(ø) sygl-1(LBS B mut)* | 4A,C,D |
| JK6391 *q869 q1116 I/hT2[qIs48](I;III)* | this work | *lst-1(ø) sygl-1(LBS alt D mut)* | S5B,E-F |
| JK6401 *q869 q828 I hT2[qIs48](I;III)* | this work | *lst-1(ø) sygl-1(ø)* | 5B; S6 |

| | | | |
|---|---|---|---|
| JK6431 *q869 q1015 I/ hT2[qIs48](I;III)* | this work | *lst-1(ø) sygl-1::V5(wt)* | 4E; 6A-C; S6B-C |
| JK6507 *q869 q1136 I/ hT2[qIs48](I;III)* | this work | *lst-1(ø) sygl-1::V5(BD mut)* | 6D-E |
| JK6508 *q1231 I* | this work | *sygl-1::1xV5(LBS B mut)* | 3B,D; S3B |
| JK6516 *q869 q1054 I/hT2[qIs48](I;III)* | this work | *lst-1(ø) sygl-1::1xV5(LBS D mut)* | 4E; 6A |
| JK6517 *q869 q1231 I/ hT2[qIs48](I;III)* | this work | *lst-1(ø) sygl-1::V5(B mut)* | 6B-E |
| JK6521 *q869 q1015 I; emb-30(tn377) III/hT2[qIs48](I;III)* | this work | *lst-1(ø) sygl-1::1xV5; emb-30* | 4F-G |
| JK6522 *q1220 I* | this work | *sygl-1::1xV5(LBS A canonical)* | S1D |
| JK6539 *q869 q1136 I/hT2[qIs48](I;III); rol-6(e187)* | this work | *lst-1(ø) sygl-1::1xV5(LBS BD mut); rol-6* | 6D-E |
| JK6566 *q1253 I* | this work | *sygl-1::1xV5(LBS CD mut)* | 3E; S4C |
| JK6567 *q869 q1054 I; emb-30(tn377) III/hT2[qIs48](I;III)* | this work | *lst-1(ø) sygl-1::1xV5(LBS D mut); emb-30* | 4F-G |
| JK6600 *q869 q1167 I* | this work | *lst-1(ø) sygl-1::V5(BCD mut)* | 6B-E; S6 |

*all 1xV5 strains also include a GGS linker (include sequence inserted)

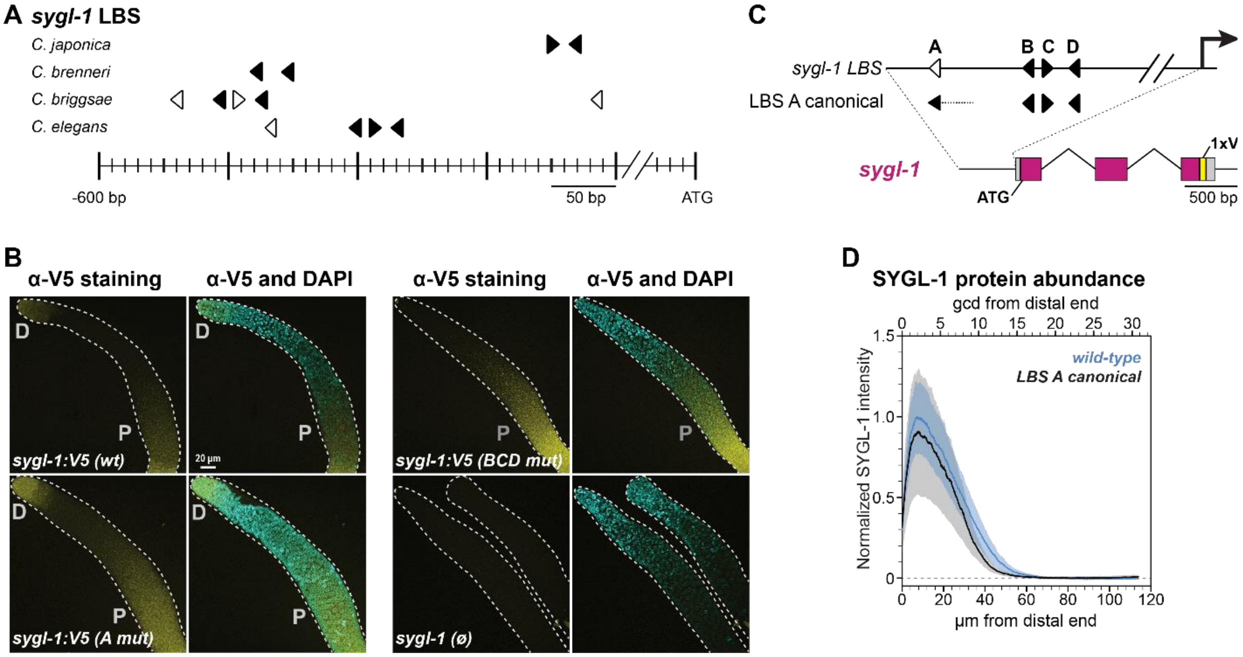


**Figure S1. Additional analyses of *sygl-1* LBS and their mutants**

**A.** LBS clusters in *sygl-1* orthologs. All canonical LBSs (filled black arrows) have the same sequence: 5' CGTGGGAA 3'; non-canonical sequences (open arrowheads) in *C. briggsae*: 5' TATGGGAA 3', 5' CATGGGAA 3', and 5' TGTGTGAA 3'. Spacing between LBSs is as follows: *C. japonica*: 10 bp; *C. brenneri*: 24 bp *C.* briggsae: from upstream to downstream 24 bp, 7 bp, 10 bp, and 252 bp; *C. elegans*: 57 bp, 7 bp, and 9 bp. **B.** Notch-independent SYGL-1 expression in the proximal gonad is not affected in LBS mutants. Lower magnification representative images of dissected gonads to show both distal and proximal SYGL-1 expression. D: distal expression; P: proximal expression. Note that wild-type proximal expression becomes brighter nearer the oocytes in the proximal germline (Lee et al. 2016); germlines lacking SYGL-1 are smaller in size and thus at this magnification more of the brighter proximal expression can be seen in germlines lacking SYGL-1. Strains and conventions are as in Fig 1F. **C-D.** Expression of SYGL-1 protein in mutants with canonical LBS A sequence. **C.** Schematic of LBS A canonical mutation. Conventions are as in Fig 1E and FigS1A. The *sygl-1* LBS A sequence was mutated from 5' AGTGGGAA 3' to 5' CGTGGGAA 3'; this mutation also replaced 23 nt downstream of LBS A (5' AAAAGGACTACTGTAGTCAATAC 3') with a PAM and *dpy-10* crRNA protospacer (5' CCGCTCGTGGTGCCTATGGTAGC 3') (dotted line). SYGL-1 was visualized by inserting a 1xV5 epitope tag at the C-terminus. **D**. The canonical LBS A mutant does not have a major effect on SYGL-1 expression. Quantification of α-V5 immunofluorescence. Pixel intensity values were extracted from sum projections made in FIJI; background N2 untagged control values (x axis) were subtracted (see Methods). Solid lines are the mean normalized intensity and shading represents SEM using n as the number of experiments. Strain genotypes in Table S1. Total gonads scored in 3 independent experiments: *wt*: 71; *LBS A canonical*: 63; untagged control: 43.

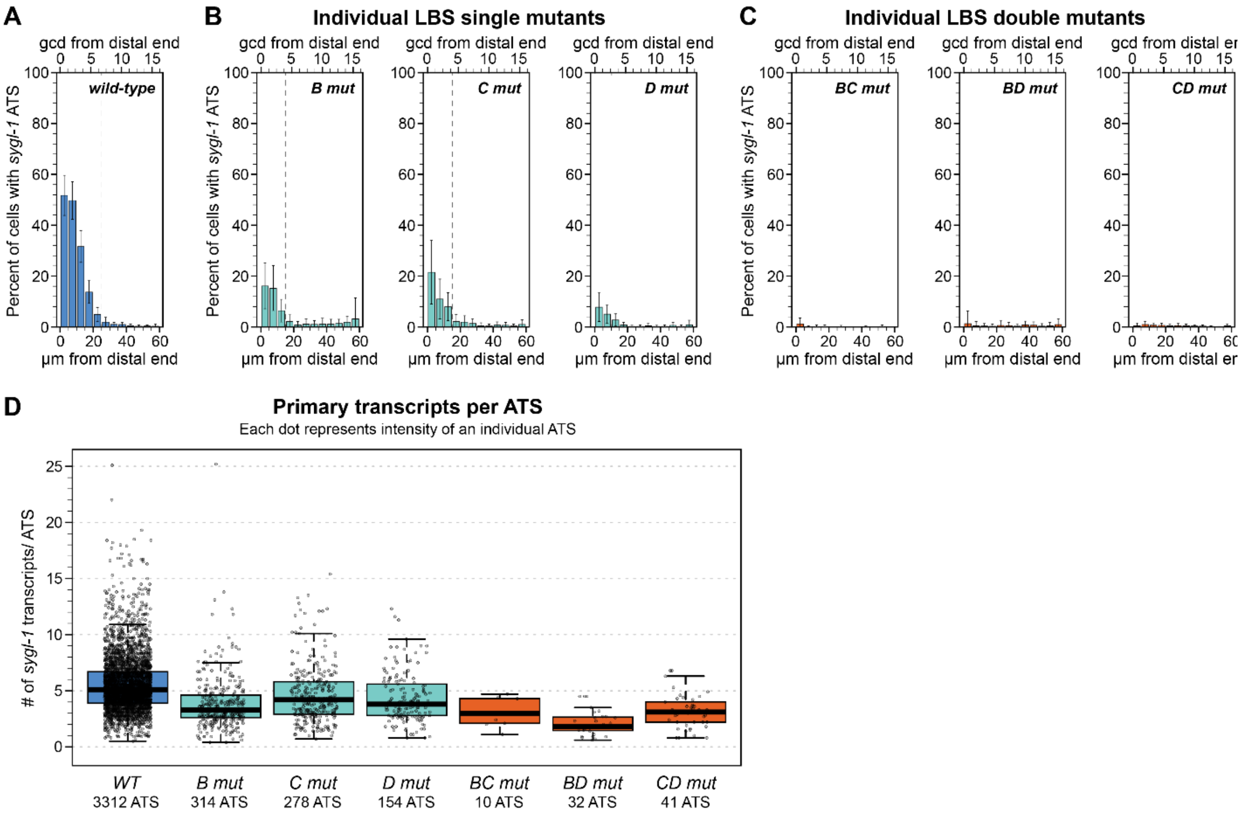


**E**

| Genotype | n | Ave # ATS/gonad | Ave # transcripts/ATS | #ATS per nucleus (%) 1 | 2 | 3 | 4 |
|---|---|---|---|---|---|---|---|
| *sygl-1(wild-type)* | **140** | **24.0** ± 8.5 | **5.6** ± 0.5 | **78** ± 8.6 | **21.0** ± 7.2 | **1.4** ± 1.5 | **0.3** ± 0.4 |
| *sygl-1(B mut)* | **50** | **6.8** ± 3.9 | **3.9** ± 0.3 | **88** ± 9.5 | **10.0** ± 7.7 | **0.8** ± 1.2 | **0.4** ± 0.6 |
| *sygl-1(C mut)* | **44** | **7.4** ± 4.6 | **4.9** ± 0.7 | **83** ± 8.8 | **14.0** ± 6.4 | **2.8** ± 2.4 | **0.3** ± 0.5 |
| *sygl-1(D mut)* | **65** | **3.1** ± 2.7 | **4.7** ± 0.9 | **89** ± 8.0 | **9.0** ± 6.1 | **0.3** ± 0.6 | **1.8** ± 2.1 |
| *sygl-1(BC mut)* | **47** | **0.3** ± 0.4 | **3.0** ± 0.2 | **100** ± 0.0 | **0.0** ± 0.0 | **0.0** ± 0.0 | **0.0** ± 0.0 |
| *sygl-1(BD mut)* | **47** | **0.9** ± 1.5 | **2.7** ± 1.1 | **92** ± 9.3 | **5.2** ± 6.0 | **1.2** ± 2.4 | **1.2** ± 2.4 |
| *sygl-1(CD mut)* | **76** | **0.7** ± 0.7 | **2.6** ± 0.9 | **96** ± 5.6 | **3.9** ± 5.6 | **0.0** ± 0.0 | **0.0** ± 0.0 |
| *sygl-1(ø)* | **51** | **0.0** ± 0.1 | **2.2** ± 0.0 | **100** ± 0.0 | **0.0** ± 0.0 | **0.0** ± 0.0 | **0.0** ± 0.0 |

**Figure S2. Transcriptional activation of individual LBS mutants is essentially equivalent**

**A-C.** ATS probability by position, shown as histogram. Percentage of nuclei expressing at least one *sygl-1* ATS as a function of distance from the distal end. Conventions as in Fig 2C. n's listed in Fig S2E. **A.** Wild-type ATS probability. Data is the same as in Fig 2C. **B.** LBS single mutant ATS probability. **C.** LBS double mutant ATS probability. **D.** Estimated number of transcripts per ATS for individual LBS mutants, shown as boxplots. Each data point is an individual ATS. Boxplot conventions as in Fig 2F; center line: median (N2: 5.1; B mut: 3.3; C mut: 4.2; *D mut*: 3.8; *BC mut*: 3.0; *BD mut*: 1.8; *CD mut*: 3.1). **E.** Summary statistics for individual genotypes that were pooled in Fig 2. Strains listed in Table S1: Numbers in table represent mean from each independent experiment plus or minus standard deviation between experimental means. n is the number of gonads scored. 9 experiments were performed and data for each genotype comes from at least 3 experiments (n experiments = N2: 9; B mut: 3; C mut: 3; *D mut*: 4; *BC mut*: 3; *BD mut*: 3; *CD mut*: 4; *sygl-1(ø)*: 3). Ave # ATS/gonad is the total number of ATS divided by the number of gonads. Average # transcripts/ATS is the mean normalized ATS intensity divided by 10 to estimate the number of nascent transcripts per ATS as in Fig 2F. #ATS per nucleus is the percent of ATS-expressing nuclei that contain the given number of ATS, as in Fig 2D.

**Figure S3**
**Lynch et al.**

A

### LBS single mutants

| Genotype | n | Percentage cytoplasmic spots with x number of mRNAs: x = 1 | x = 2 | x = 3 | x = 4+ |
|---|---|---|---|---|---|
| *sygl-1(wild-type)* | **140** | **61** ± 4 | **30** ± 4 | **6.2** ± 0.7 | **2.3** ± 0.3 |
| *sygl-1(B mut)* | **50** | **62** ± 4 | **33** ± 4 | **4.6** ± 0.3 | **1.0** ± 0.4 |
| *sygl-1(C mut)* | **44** | **61** ± 4 | **32** ± 5 | **5.2** ± 0.5 | **1.5** ± 0.2 |
| *sygl-1(D mut)* | **65** | **59** ± 2 | **35** ± 1 | **5.0** ± 0.8 | **1.1** ± 0.4 |
| *sygl-1(ø)* | **51** | **73** ± 4 | **20** ± 5 | **5.8** ± 0.7 | **2.0** ± 0.4 |

n: number of gonads scored
x: number of mRNAs estimated per cytoplasmic spot

### LBS double mutants

| Genotype | n | Percentage cytoplasmic spots with x number of mRNAs: x = 1 | x = 2 | x = 3 | x = 4+ |
|---|---|---|---|---|---|
| *sygl-1(wild-type)* | **140** | **61** ± 4 | **30** ± 4 | **6.2** ± 0.7 | **2.3** ± 0.3 |
| *sygl-1(BC mut)* | **47** | **59** ± 7 | **37** ± 8 | **4.6** ± 1.9 | **0.3** ± 0.5 |
| *sygl-1(BD mut)* | **47** | **60** ± 8 | **37** ± 9 | **2.9** ± 0.9 | **0.6** ± 0.5 |
| *sygl-1(CD mut)* | **76** | **57** ± 8 | **39** ± 9 | **3.1** ± 1.3 | **0.6** ± 0.4 |
| *sygl-1(ø)* | **51** | **73** ± 4 | **20** ± 5 | **5.8** ± 0.7 | **2.0** ± 0.4 |

n: number of gonads scored
x: number of mRNAs estimated per cytoplasmic spot

B α-V5 staining α-V5 and DAPI staining

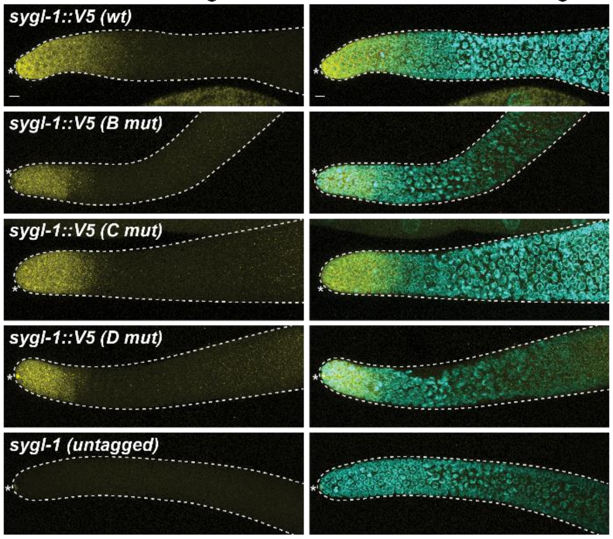


C α-V5 staining α-V5 and DAPI staining

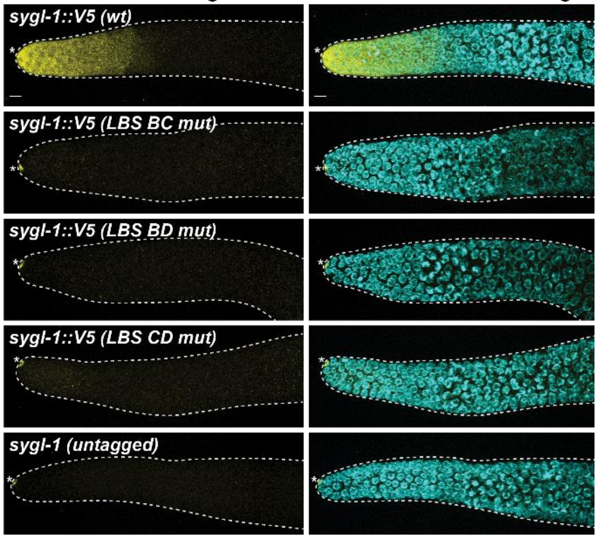


**Figure S3. Characterization of LBS single mutant mRNA and protein data**
**A.** Percent of cytoplasmic spots containing 1, 2, 3, or 4+ mRNA as a ratio of the total number of cytoplasmic spots detected. Numbers are averages per replicate plus or minus the standard deviation between replicates. n: number of gonads analyzed, from 3-9 independent experiments. **B.** Representative images of SYGL-1 protein inLBS single mut gonads. Images were selected from the same experiment, were adjusted with the same contrast values in FIJI, and are maximum z projections. Dashed gray line outlines the gonad and asterisk marks the niche cell body. V5 signal stains the niche cell body nonspecifically. Scale bar: 5 µm. **C.** Representative images of SYGL-1 protein in LBS double mut gonads; conventions as in Fig S3B.

**A** **Average number of MATLAB-detected nuclei per bin**

| | Bin number | | | | | | | | | | | |
|---|---|---|---|---|---|---|---|---|---|---|---|---|
| **Genotype** | **1** | **2** | **3** | **4** | **5** | **6** | **7** | **8** | **9** | **10** | **11** | **12** |
| *sygl-1(wild-type)* | 8.5 | 12.5 | 15.3 | 17.3 | 19.3 | 20.0 | 21.5 | 22.1 | 22.3 | 22.1 | 20.3 | 9.1 |
| *sygl-1(B mut)* | 7.9 | 11.9 | 13.9 | 15.6 | 16.4 | 16.6 | 17.3 | 18.3 | 17.6 | 17.3 | 16.5 | 8.9 |
| *sygl-1(C mut)* | 7.8 | 12.0 | 14.3 | 16.5 | 16.6 | 18.0 | 18.1 | 19.0 | 19.5 | 18.7 | 19.1 | 9.0 |
| *sygl-1(D mut)* | 7.8 | 11.9 | 14.6 | 17.5 | 18.9 | 20.1 | 20.4 | 21.3 | 20.1 | 20.0 | 18.1 | 8.9 |
| *sygl-1(BC mut)* | 8.8 | 13.5 | 16.4 | 17.8 | 19.2 | 19.0 | 19.4 | 19.7 | 18.6 | 17.6 | 15.5 | 7.4 |
| *sygl-1(BD mut)* | 8.4 | 13.7 | 16.5 | 18.6 | 19.4 | 20.2 | 20.0 | 19.7 | 19.5 | 17.7 | 16.3 | 8.1 |
| *sygl-1(CD mut)* | 8.4 | 12.5 | 16.0 | 16.7 | 17.9 | 18.7 | 19.0 | 18.4 | 17.7 | 17.3 | 15.8 | 7.8 |
| *sygl-1(ø)* | 9.0 | 13.6 | 15.7 | 17.8 | 18.4 | 18.1 | 18.1 | 16.9 | 16.6 | 14.8 | 13.9 | 6.9 |

**B** **LBS effects on mRNA distribution: empty cells vs. cells containing *sygl-1* mRNA**

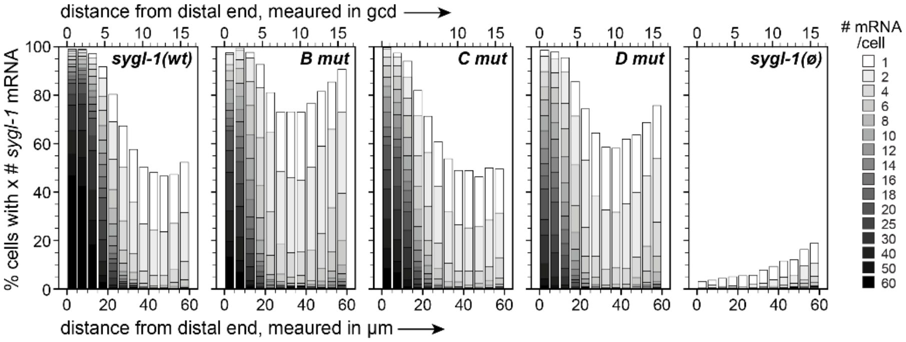


**Figure S4. Analyzing the cell-to-cell distribution of *sygl-1* mRNA in LBS mutants**

**A.** Average number of nuclei per bin (region from the distal end). MATLAB reconstructs the gonad in 3D and assigns each nucleus center with an x coordinate (Lee et al., 2016). Bin 1 corresponds to 0-5 µm from the distal end, bin 2 to 5-10 µm, and so on. The average nuclei/bin numbers were taken from the smFISH data set; see Fig. S2E for the total number of gonads scored for each genotype. Strain genotypes in Table S1. **B.** mRNA probability by position, shown as histograms. Conventions as in Fig. 5D, except that #mRNA/cell key has 15 colors rather than three. The maximum numbers of mRNA/cell are as follows: *wt*: 146; *B mut*: 90; *C mut*: 111; *D mut*: 86; *sygl-1(ø)*: 3.

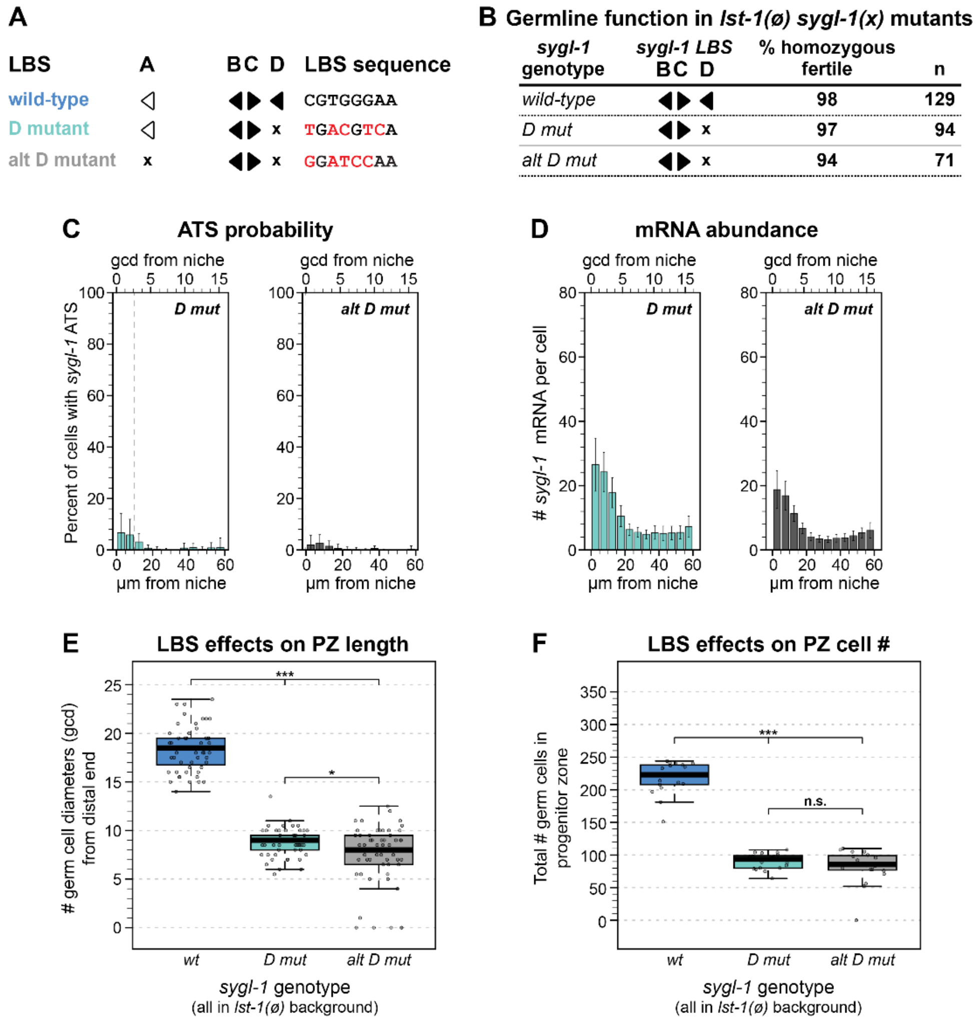


**Figure S5. An alternatively mutated LBS sequence, *alt D mut*, is homozygous fertile like other LBS singles but has marginal changes in *sygl-1* dose**

**A.** Schematic of two distinct 5bp LBS D mutations generated by Cas9 editing. Conventions as in Fig 1E. Wild-type: 5' CGTGGGAA 3'; *D mut*: 5' TGACGTCA 3'; *altD mut*: 5' GGATCCAA 3' (differences from wild-type motif underlined). Unlike *D mut*, *alt D mut* mutates all three of the central guanines. In *alt D mut*, we mutated the middle G to a C: typically, the middle G is recognized as the most degenerate of the trio of guanines, but there is also *in vitro* evidence that suggests a mutation to C is particularly deleterious for CSL binding (Torella et al. 2014, Friedmann and Kovall 2010, others). **B-F.** *alt D mut* and *D mut* were directly compared. As in the rest of the paper, effects on molecular quantitation was scored in *lst-1(+)* (Fig C-D) while effects on phenotype were scored in *lst-1(ø)* (Fig B, E-F). See Table S1 for strain genotypes. **B.** Germline fertility was scored by presence of embryos in young adults, as in Fig. 4A. **C-D.** alt *D mut* (n = 56 gonads) was directly compared to *D mut* (n = 39 gonads) in 2 smFISH experiments. **C.** ATS probability as a function of distance from distal end. Conventions are as in Fig 2C. **D.** Number of mRNA per cell as a function of distance from the distal end. Conventions are as in Fig 3A. **E.** PZ length shown as boxplots, BoxPlotR conventions; center line: median. PZ observations from each gonad were fitted to a linear mixed effects model and Tukey's post-hoc test was used to make pairwise comparisons between genotypes. ***: $p < 0.0001$; *: $p = 0.009$. Total gonads scored from two independent experiments: *wt:* 55; *altD mut*: 57; *D mut*: 57. **F.** PZ cell number shown as boxplots, BoxplotR conventions. Automatic counts in DAO-5-stained gonads. ***: $p < 0.0001$; n.s.: $p = 0.43$. Total gonads scored: *wt:* 18; *altD mut*: 20, *D mut*: 20 from one experiment.

**Figure S6**
**Lynch et al.**

**A** Two versions of *sygl-1* null

| | LBS motifs intact? | *sygl-1* CDS present? | V5 tag on *sygl-1* C-term? |
|---|---|---|---|
| ***sygl-1(ø)*** | yes | no | no |
| ***sygl-1::V5(BCD mut)*** | no | yes | yes |

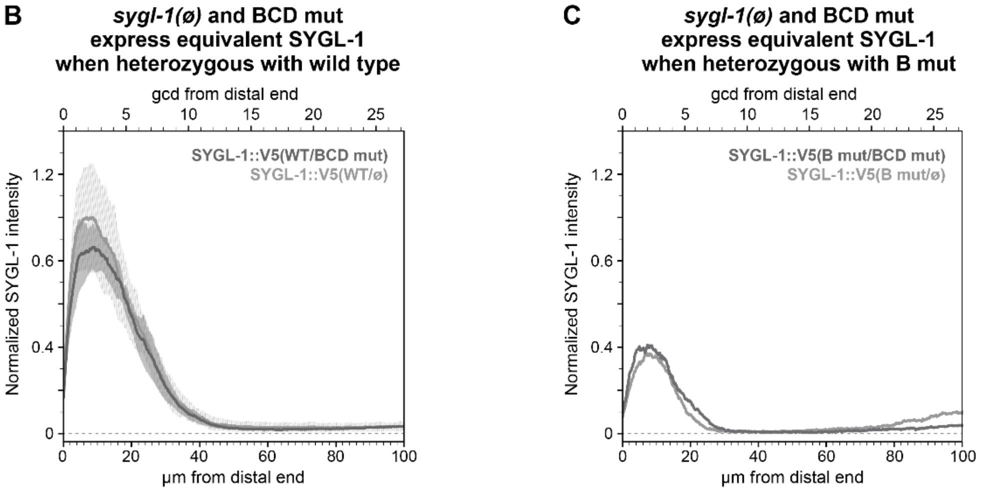


**Figure S6. Using *sygl-1(null)* strains in heterozygotes**
**A.** Both *sygl-1(ø)* (Fig 2A) and *sygl-1(BCD mut)* (Fig 1E) are null in the distal germline but differ in a few ways that made *BCD mut* a more suitable null allele for exploring LBS number and SYGL-1 dose. In *sygl-1(ø)*, Cas9 editing removed the *sygl-1* ORF from the start of the first exon to the end of the last exon, but left UTRs and LBS intact. There is no V5 tag on *sygl-1(ø)*. In *BCD mut*, the *sygl-1* ORF is intact and V5-tagged, but all three LBSs are mutated. Neither allele expresses detectable SYGL-1 protein in the distal germline (Fig 1F-G). **B-C.** *wt/null* or *B mut/null* heterozygotes produce similar results regardless of *sygl-1* null allele used. Quantification of α-V5 immunofluorescence as a function of distance from the distal end, conventions as in Fig 3C. **B.** Total gonads scored: *wt/ø*: 24; *wt/BCD mut*: 30, from 2 independent experiments each (*wt/ø* and *wt/BCD mut* immunostaining not done in parallel). **C.** Total gonads scored: 10 each for *B mut/ø* and *B mut/BCD mut*, immunostaining compared in parallel from one experiment.

**Figure S7**
**Lynch et al.**

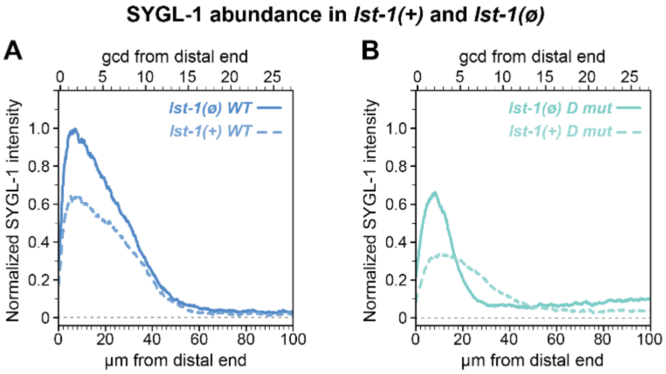


**Figure S7. Absence of *lst-1* increases SYGL-1 abundance**

**A-B.** Quantification of α-V5 immunofluorescence. All four strains were assayed in parallel but separated into two graphs for clarity. 24 gonads/genotype scored in 2 replicates. The mechanism by which *lst-1* removal changes SYGL-1 abundance is not understood. However, understanding that SYGL-1 abundance is not the same in *lst-1(+)* and *lst-1(ø)* is relevant to this paper. **A.** *sygl-1::V5(wt)* abundance compared in *lst-1(+)* and *lst-1(ø)* backgrounds. **B.** *sygl-1::V5(D mut)* abundance compared in *lst-1(+)* and *lst-1(ø)* backgrounds.
